# Absolute Quantification of Lysosomal Proteins by Multiple Reaction Monitoring Mass Spectrometry and QconCAT Protein Standards

**DOI:** 10.1101/2025.01.09.632238

**Authors:** Peter Robert Mosen, Biswajit Moharana, Sofía Fajardo-Callejon, Edgar Kaade, Norbert Rösel, Maryam Omidi, Roman Sakson, Thomas Ruppert, Sandra Pohl, Volkmar Gieselmann, Dominic Winter

## Abstract

**ABSTRACT:** Lysosomes are membrane-enclosed organelles that play a crucial role in the degradation of intra- and extracellular substrates and the regulation of metabolic signaling. These functions are carried out by a variety of proteins, of which > 150 are currently known to be located in the lysosomal lumen or to be embedded in its membrane. These proteins are typically low abundant, necessitating organelle enrichment experiments to enable their investigation by unbiased mass spectrometry-based proteomics analyses. Enrichment strategies have been applied in a plethora of studies to gain a deeper understanding of lysosomal composition and relative changes of lysosomal proteome abundance in a variety of pathological conditions. Such experiments are restricted, however, to selected cell lines and tissues and do not allow a direct analysis of the lysosomal proteome from whole cell or tissue lysates. Furthermore, they do not provide absolute quantities. We developed a multiple reaction monitoring mass spectrometry-based assay for the highly sensitive and reproducible absolute quantification of 143 mouse lysosomal proteins from any type of sample following the QconCAT strategy. We applied our approach to the investigation of mouse embryonic fibroblast whole cell lysates and lysosome-enriched fractions, providing absolute copy numbers per cell/lysosome for lysosomal hydrolases and membrane proteins. These data reveal a dynamic range of more than three orders of magnitude in protein expression and strong differences in the subcellular distribution of individual lysosomal proteins. Furthermore, we applied our strategy to the investigation of four primary cell types (macrophages, lung fibroblasts, osteoblasts, and osteoclasts), demonstrating pathway-specific heterogeneity of individual lysosomal protein classes and indicating protein-specific post-transcriptional regulation of expression levels.

## Introduction

Lysosomes are membrane-enclosed organelles which are present in almost all eukaryotic cells, and present both the cell’s main degradative compartment and their center for metabolic signaling [1, 2]. The lysosomal lumen is characterized by an acidic pH (pH 4.5-5.5) and contains a variety of hydrolases which catalyze the degradation of most biological macromolecules (nucleic acids, proteins, lipids, and carbohydrates), ranging from single molecules to large structures such as protein complexes, organelles, and whole cells [3–6]. These substrates are delivered to lysosomes through different routes, which can be broadly categorized in the endocytic pathway, for extracellular material and plasma membrane-localized proteins, and the autophagic pathway, for intracellular material [1, 7].

While > 340 proteins have been described to be related to correct lysosomal function irrespective of their subcellular localization so-far, ∼150 were shown to be lysosomal-resident, i.e. to be localized either in its lumen or to contain at least one domain spanning its membrane [8]. Broadly, these proteins can be classified into hydrolases and their co-factors facilitating substrate degradation, membrane proteins providing stability and protection from proteolytic digestion, and transporters/exchangers with their accompanying subunits to recycle generated nutrients and to regulate ion homeostasis [8, 9].

Mutations resulting in the alteration/loss of function of lysosomal proteins result in a class of ∼70 rare inherited diseases (cumulative incidence of 1 in 5000 live births), termed lysosomal storage disorders (LSDs) [10]. LSDs are characterized by an accumulation of lysosomal substrates, which gradually incapacitates organellar function, initiating a complex pathological cascade that affects a variety of cellular processes [11]. While LSDs and their detrimental effects on cellular function have been known since the 1950s, there is a growing body of evidence that alterations of lysosomal proteins are also causative for more common conditions such as neurodegenerative diseases (NDs, e.g. Parkinson’s disease [12–17] and various types of cancer [18–24].

Furthermore, the view of lysosomes as being static and unregulated, which prevailed for decades, is currently replaced by a picture of highly motile and dynamic organelles [25–28] that are actively responding to alterations in cellular homeostasis [2, 29] and play a key role in the regulation of a large variety of cellular processes [25, 30–36]. Given these highly diverse roles the lysosome is involved in, it is not surprising that recent studies demonstrated lysosomal heterogeneity both between different types of cells [37, 38] and within the same cell [27] which is also in line with the observation that lysosome-related diseases present with cell-/tissue-specific phenotypes [10].

Distinct lysosomal proteins and lysosomal morphology have been thoroughly investigated in a large variety of cells, tissues, and disease-contexts by western blotting, immunohistochemistry and electron microscopy [39–41]. While this resulted in a wealth of data facilitating a general understanding of disease mechanisms and their immediate effects on cellular (patho-) physiology, there remains a significant gap in comprehending the broader response of the lysosomal proteome to cellular homeostasis, disease-causing mutations, and therapeutic interventions.

For studies addressing the complete set of lysosomal proteins – the lysosomal proteome – mass spectrometry (MS)-based proteomics presents an ideal tool [37, 38, 42]. So-far, however, MS studies of lysosomes have been mainly restricted to immortalized cell lines and few types of tissue [8, 43], which are compatible with the enrichment of lysosomes. This is due the low abundance of lysosomal proteins (it is currently estimated that they comprise ∼0.2% of cellular biomass [44], which results in their incomplete coverage between individual samples and insufficient analytical performance in whole cell proteomics experiments [37]. Furthermore, MS-based proteomics of lysosomal proteins has been restricted to relative comparisons between similar sample types. This is due to sample matrix effects on the signal intensity of individual peptides, as they result in intensity differences for the same peptide between samples of differing complexity. Due to these limitations, our current knowledge of the composition of the lysosomal proteome, and its response to pathological conditions, is largely limited to relative comparisons within the same tissue and cell culture-based systems [37, 42, 45].

An approach which enables the highly sensitive and reproducible quantification of proteins in virtually any type of samples by MS is targeted proteomics by parallel/multiple reaction monitoring (PRM/MRM) [46, 47]. Compared to unbiased whole proteome analyses, PRM/MRM analyses are based on the manual selection of surrogate peptides representing distinct proteins of interest, which are subsequently exclusively targeted by the mass spectrometer [48]. In combination with stable isotope-labeled standard peptides, this further allows for the absolute quantification of proteins [49]. For example, targeted proteomics has been demonstrated to enable the highly reproducible quantification of even lowest protein amounts from cell lysates [50, 51], as well as body fluids such as human plasma/serum [52].

Typically, stable isotope-labeled standard peptides are generated by chemical synthesis. While this is undoubtedly a straightforward approach, its use on large numbers of peptides is restricted by the high costs of peptides, as they have to be individually purified and absolutely quantified. The QconCAT strategy, which is based on engineered artificial standard proteins (so-called Quantification conCATemers, QconCATs [53, 54]), presents an attractive alternative for the affordable generation of large numbers of absolutely quantified internal standards. In a recent benchmark study, this approach was utilized for the absolute quantification of ∼2000 peptides in yeast [55].

In the current study, we developed a strategy for the MRM-based absolute quantification of 143 mouse lysosomal proteins utilizing QconCAT-derived absolutely quantified stable isotope-labeled standard peptides. We provide protein copy number information per cell/organelle for mouse embryonic fibroblasts (MEFs), lung fibroblasts, osteoclasts, osteoblasts, and macrophages. We identify differences in lysosomal protein expression and subcellular distribution, and correlation of protein copy numbers and transcript levels reveal cell type-specific posttranscriptional regulation of lysosomal protein amounts.

## Material and Methods

### Peptide selection and construction of QconCAT protein standards

Based on analyses of in-house and published datasets [56–59] ≥ 2 peptides/lysosomal protein were manually selected from a pool of > 6300 unique tryptic peptides considering the following factors: a) proteotypicity, b) signal intensity, c) possibility of post-translational or chemical modifications (PTM), d) missed cleavage sites, e) proximity to proline, acidic amino acids or protein termini, and f) cleavage between two basic residues. If more than three peptides per lysosomal protein were available, the three most intense ones were chosen. Selected lysosomal protein-derived peptides were grouped based on their abundance (averaged across individual peptides) in large scale proteomics datasets of brain and liver [56, 57, 59] into 12 groups (from highest to lowest intensities in QconCAT 1 to QconCAT 12) with all peptides representing a given lysosomal protein on the same QconCAT. Subsequently, the order of peptides in each QconCAT was optimized in-silico via PeptideCutter [60] to achieve maximum cleavage efficiency. To enable absolute quantification of individual QconCATs, a minimally permutated peptide analog (MIPA, [61]) peptide was placed at the N-terminus, and a 6x His-tag was placed at the C-terminus for Ni^2+^affinity purification, of each QconCAT.

### Plasmid design and expression of stable isotope-labeled QconCATs

Designed QconCAT protein sequences were reverse-translated, codon-optimized, generated by gene synthesis, and cloned into pET-28a (+) expression vectors (69864-3, EMD Biosciences). To enhance bacterial expression efficiency of lysosomal QconCAT 12, it was further fused with maltose-binding protein (MBP, [62]) and the MBP-QconCAT 12 fusion construct was sub-cloned into the PBAD-his6- prkA-pACYC184 (pRKA) expression vector [63]. pRKA was a gift from Ichiro Matsumura (Addgene plasmid # 41041; http://n2t.net/addgene:41041; RRID: Addgene_41041). Individual QconCAT plasmids were transformed into *E. coli* BL21 DE3 cells and stable isotope-labeled protein constructs were expressed and purified as described elsewhere [53]. Briefly, bacteria were grown in M9 minimal medium (including kanamycin for QconCATs 1-11 and ampicillin for QconCAT 12) [53], containing ^13^C_615_N_4_ arginine (CNLM-539-H-0.05) and ^13^C_615_N_2_ lysine (CNLM-291-H-0.05, both Cambridge Isotope Laboratories) until an optical density (OD) of 0.6-0.8 was reached. Subsequently QconCAT protein expression was induced by addition of 1 mM (final concentration) isopropyl β-D-1-thiogalactopyranoside (IPTG, Thermo Fisher Scientific) and bacterial cultures incubated without agitation at RT for 4-6 hrs until reaching a final OD of 0.8-1. Bacteria were harvested by centrifugation at 8000x g, 4 °C for 10 min, the supernatant discarded, and cell pellets were stored at -20 °C. For cell lysis, bacteria were resuspended in 2.5 ml of B-PER bacterial lysis buffer (Thermo Fisher Scientific) per 1 g of cell pellet, centrifuged at 6000x g, 4 °C for 10 min, the supernatant was discarded, and the pellet resuspended in 2.5 ml fresh B-PER. Lysozyme (0.2 mg/ml, Sigma Aldrich), protease inhibitor cocktail (1x c0mplete EDTA-free, Roche) and DNaseI (300 U, Thermo Fisher) were added (all final concentrations) and the bacterial lysate was incubated on a horizontal shaker at RT for 10 min. Subsequently, 15 ml of a 1:10 B-PER dilution (in water) was added and the solution was vortexed and centrifuged at 15,000x g, 4 °C for 15 min. The pellet containing bacterial inclusion bodies was washed twice with 20 ml of a 1:10 B-PER dilution and stored at -20 °C until further use. Inclusion bodies were solubilized in 1% sodium dodecyl sulfate (SDS) or 6 M guanidinium chloride (GCl) in 500 mM NaCl, and 20 mM NaH_2_PO_4_/Na_2_HPO_4_ (pH 7.4). Samples were diluted to a final volume of 20 ml with 0.1% SDS or 6 M GCl, centrifuged at 15,000x g, RT for 15 min and purified by pre-equilibrated nickel-charged 1 ml His- trap columns (17-5247-01, GE Healthcare). Bound proteins were eluted using 500 mM NaCl, 20 mM NaH_2_PO_4_/Na_2_HPO_4_, 500 mM imidazole, and 50 mM EDTA (pH 7.4) in 0.1% SDS or 6 M GCl and stored at -20 °C.

### Cell culture and lysosome enrichment

All cell culture experiments were performed under sterile conditions and cells cultured at 37 °C, 100% humidity, and 5% CO_2_. Mouse embryonic fibroblasts (MEFs) were seeded at a density of 1.5×10^6^ cells and cultured for 72 hrs using DMEM supplemented with 10% fetal bovine serum (FBS), 100 mU/ml penicillin, 100 µg/ml streptomycin, and 2 mM glutamine (all Thermo Fisher Scientific). For macrophage, osteoclast and osteoblasts differentiation, the bone marrow was flushed out of the femora from six wild-type mice (C57BL/6 mouse) per cell type at the age of 12 weeks with α-minimal essential medium (α- MEM) containing 10% FBS. The care and use of mice complied with all relevant local animal welfare laws, guidelines and policies (ORG-1091). Animal experiments were approved by the animal facility of the University Medical Center Hamburg-Eppendorf and by the Behörde für Gesundheit und Verbraucherschutz. Wild-type C57BL/6 mice were housed in a pathogen-free facility at the University Medical Center Hamburg-Eppendorf, with a 12-hour light/dark cycle, 45% to 65% relative humidity, and an ambient temperature of 20°C to 24°C. Experimental procedures were conducted following institutional guidelines. Extracted cells were plated at a density of 12.5×10^6^ cells per well, incubated for 24 hrs, the media was removed, and dead cells were washed off. Adherent cells were further cultured in α-MEM containing 10% FBS and 10 nM 1,25-dihydroxyvitamin-D3 (Sigma Aldrich). Beginning at day 4 after seeding, M-Csf (recomb. murine, 315-02, Peprotech) and receptor activator of NF-κB ligand (recomb. murine *E.Coli* derived, 315-11, Peprotech) were added to a final concentration of 20 ng/ml and 40 ng/ml, respectively, and cells were cultured for 7 days to generate osteoclasts. Primary macrophages were generated by the same method without addition of Rankl (NF-κB ligand). For osteoblast differentiation, bone marrow cells were cultured for 10 days in α-MEM containing 10% fetal bovine serum, 25 µg/ml L-ascorbic acid (Sigma Aldrich), and 5 mM ß-glycerophosphate (Sigma Aldrich). For lung fibroblasts, lungs of six P12 wild-type mice were dissected and enzymatically digested in 1x phosphate-buffered saline (PBS) containing collagenase type III (150 U/ml, PAN Biotech) and dispase I (2.5 U/ml, Sigma-Aldrich). After incubation at 180 rpm, 37 °C for 3 hrs, strained cells were seeded (12.5×10^6^ cells per well) and cultured for 7 days. For harvesting, cells were washed with ice-cold 1x PBS, scraped in 500 µl ice-cold PBS, and centrifuged at 1000x g, 4 °C for 10 min. Supernatants were discarded and the cell pellet was stored at -80 °C. Lysosomes were enriched from MEFs using superparamagnetic iron oxide nanoparticles (SPIONs) from two 10 cm plates with a seeding density of 3×10^6^ cells as described elsewhere [64]. Briefly, cells were incubated with SPIONs for 24 hrs, followed by 24 hrs chase, harvested by scraping, lysed using a dounce homogenizer, and lysosomes were enriched from the postnuclear supernatant using LS-columns and a magnetic stand (Miltenyi Biotec). Lysosome-enriched fractions were concentrated by centrifugation at 20,000x g, 4° C for 30 min and the supernatant discarded. For the determination of cellular protein content of MEFs, cells were harvested by trypsinization, cell numbers determined using a Neubauer chamber, and the protein content of defined amounts of cells determined using the DC protein assay (Bio-Rad).

### Proteolytic digestion of QconCATs

To evaluate the performance of different digestion strategies on peptide release from QconCATs, proteins eluted in 0.1% SDS were digested by single-pot, solid-phase-enhanced sample-preparation (SP3), RapiGest, or in gel digestion (IGD), and such eluted in 6 M GCl in solution using GCl or Filter Aided Sample Preparation (FASP), as described elsewhere [65–69] with minor modifications. In the following, all approaches are briefly outlined. For reduction/alkylation reagents final concentrations are indicated. For SP3, 10 µg of protein were adjusted to a total volume of 10 µl with 1% SDS, 50 mM HEPES (pH 8) in PCR-tubes. Proteins were reduced with 5 mM dithiothreitol (DTT) at 56 °C for 45 min, alkylated with 20 mM acrylamide (AA) at RT for 30 min, and the reaction was quenched with 5 mM DTT at RT for 10 min. SP3 beads (1:1 mixture of Sera-Mag SpeedBeads 45152105050250 and 65152105050250, GE Healthcare) were added at a 10:1 (w/w) bead to protein ratio and protein binding was induced by addition of ethanol (final concentration: 80%) and incubation at 1000 rpm, RT for 20 min on a thermomixer. Using a magnetic rack, the supernatant was discarded, the particle-bound proteins were washed (2x with 150 µl 80% ethanol and 1x with 100 µl 100% ACN), and the supernatant removed. Subsequently, proteins were proteolytically digested by addition of 50 µl 100 mM ammonium bicarbonate (ABC) solution containing trypsin at a 1:50 (w/w) protease-to-protein ratio and incubation in a thermomixer at 1000 rpm, 37 °C overnight. The next day, beads were immobilized using the magnetic rack and the peptide-containing supernatants transferred to fresh tubes. For RapiGest digestion, 0.25 µg of protein were resuspended in 1% RapiGest (Waters), diluted 1:4 with 0.133 M ABC to a final volume of 20 µl, reduced with 5 mM DTT at 56 °C for 45 min, alkylated with 20 mM AA at RT for 30 min, and the reaction was quenched with 5 mM DTT for 10 min at RT [66, 70]. Protein digestion was performed at 37 °C overnight using trypsin at a protease-to-protein ratio of 1:5 (w/w) in a final volume of 30 µl, and a final concentration of 0.1% RapiGest in 100 mM ABC. The next day, RapiGest was hydrolyzed by addition of 1% (final concentration) trifluoroacetic acid (TFA) and incubation at 800 rpm, 37 °C for 45 min. RapiGest hydrolysis products were precipitated by centrifugation at 14,000x g, RT for 15 min and peptide-containing supernatants were transferred to fresh tubes. For in gel digestion, 75 µg of protein were combined with 4x Laemmli loading buffer (8% SDS (w/v), 10% β-mercaptoethanol (BME), 10% (v/v), 40% glycerol (v/v), 0.04% bromophenol blue (w/v), 0.5 M Tris-HCl, pH: 6.8) to reach a final concentration of 1x [71]. Samples were reduced at 95 °C for 10 min, alkylated with 20 mM AA at RT for 30 min, and the reaction was quenched with 5 mM DTT at RT for 10 min. A sample volume corresponding to 60 µg protein was distributed across 9 lanes of a 10% polyacrylamide gel (∼6.7 µg/lane), SDS-PAGE was performed at a constant voltage of 120 V for 90 min, the gels washed with milliQ water, and stained overnight using Coomassie brilliant blue (Thermo Fisher Scientific). The next day, gels were washed three times with milliQ water, the QconCAT bands excised and cut to small pieces, pooled, and transferred to a 1.5 ml tube. Gel pieces were de-stained using 30% ACN 70 mM ABC, dehydrated with ACN, and dried using a vacuum centrifuge. Subsequently, gel pieces were rehydrated with 100 mM ABC containing trypsin at a protease-to-protein ratio of 1:25 (w/w), digested overnight at 37 °C and the peptides recovered the next day by successive incubation of the gel pieces with 50% ACN 0.1% TFA, 100 mM ABC, and 50% ACN 50 mM ABC. Recovered peptide-containing supernatants were pooled and dried using a vacuum centrifuge. For FASP, all incubation steps were followed by centrifugation of the filter unit at 14,000x g, RT for 15 min and discarding of the flow through unless stated otherwise. For each sample, 20 µg of protein was brought to a final volume of 300 µl with 6 M GCl, 100 mM Tris-HCl, pH 7.5. Samples were loaded on equilibrated centrifugal filter units (Microcon-30kDa, MRCF0R030, Merck) and samples were concentrated at 14,000x g, RT for 25 min. Subsequently, the filter units were washed twice with 100 µl of 6 M GCl, 100 mM Tris-HCl. Proteins were reduced with 10 mM BME at 56 °C for 45 min, alkylated with 10 mM chloroacetamide at RT for 30 min and centrifugal filter units washed three times with 100 µl Tris-HCl (pH 7.5) and twice with 100 µl 100 mM ABC (pH 8). Samples were digested by addition of trypsin in 50 µl 100 mM ABC at a protease-to-protein ratio of 1:100 (w/w) and incubation at 37 °C overnight in a wet chamber. Peptides were recovered by centrifugation and such retained on the filter membrane eluted by an additional centrifugation step with 50 µl 500 mM NaCl. For GCl in solution digestion, an equimolar mix of QconCAT proteins (1 µg per QconCAT protein) was prepared and adjusted to 30 µl of 5.5 M GCl in 100 mM HEPES (pH 7.6). Proteins were reduced with 10 mM BME at 56 °C for 45 min, alkylated with 20 mM chloroacetamide at RT for 30 min, and the reaction was quenched with 10 mM BME at RT for 10 min. For proteolytic digestion the GCl concentration was further reduced to 1 M, trypsin added at a 1:5 (w/w) protease-to-protein ratio, and incubated overnight at 37 °C.

### Proteolytic digestion of cell pellets and lysosome-enriched fractions

Cell/lysosome pellets were resuspended in 4% SDS, 0.1 M HEPES pH 8.0 at a pellet to buffer ratio of 1:10 (v/v), heated at 95 °C for 10 min, and sonicated three times for 30 s at 60% duty cycle and an output of 6 with ultrasonic processor (MSK-USP-3, 300W) on ice until complete solubilization of pellet was observed. Lysates were incubated again at 95 °C for 5 min, centrifuged at 20,000x g, RT for 30 min, and the clear supernatants transferred to fresh tubes followed by determination of protein concentrations using the DC protein assay. Samples were stored at -80 °C until further use. For each sample, 100/250 µg of protein was precipitated with methanol/chloroform as described elsewhere [72], and pellets were solubilized in 0.2% RapiGest, 200 mM ABC (pH 8) by incubation at 400 rpm, 95 °C for 15 min, ultrasonication for 30 min at RT in a water bath, and pre-digestion with trypsin at 37 °C for 45 min at a protease-to-protein ratio of 1:500. Subsequently samples were diluted to 50% of the final digestion volume, proteins were reduced using 5 mM DTT at 56 °C for 45 min, alkylated with 20 mM AA at RT for 30 min, and the reaction was quenched with 5mM DTT (all final concentrations) [73]. Samples were adjusted to 0.1% RapiGest at protein concentration of ≥ 1µg/µl [70] and trypsin was added at a protease-to-protein ratio of 1:50. Proteolytic digestion was performed in a thermomixer at 500 rpm, 37 °C overnight. Samples were acidified with 1% TFA (final concentration), incubated at 37

°C for 45 min, centrifuged at 14,000x g, RT for 15 min, and the supernatants transferred to a fresh tube.

### Peptide desalting and quantification

Peptides from digests with inputs of > 75 µg were desalted with OASIS HLB cartridges (1cc 10mg, Waters) and such with < 75 µg with C_18_ STAGE tips (Empore discs, # 2215, 3M) as described elsewhere [74]. Briefly, tips were activated with MeOH and cartridges/tips washed with and 80% ACN, 0.5% acetic acid (AA) and 0.5% AA. Samples were acidified with 0.5% AA (final concentration), loaded to the cartridge/tip, washed with 0.5% AA, and eluted with 80% ACN, 0.5% AA. Eluate fractions were dried using a vacuum centrifuge, resuspended in 5% ACN, and peptide amounts were determined using a fluorometric peptide assay (23290, Thermo Fisher Scientific).

### Liquid chromatography tandem mass spectrometry data acquisition

The following liquid chromatography tandem mass spectrometry (LC-MSMS) setups were used: an EASY-nLC 1000 UHPLC coupled to an LTQ Orbitrap Velos (both Thermo Fisher Scientific), a Dionex Ultimate 3000 UHPLC coupled to an Orbitrap Fusion Lumos (both Thermo Fisher Scientific), a nano ACQUITY UPLC system (Waters) coupled to a QTRAP 5500 (Sciex) and a Dionex Ultimate 3000 UHPLC coupled to a QTrap 6500+ (Sciex). If not indicated otherwise, spray tips were produced in-house from fused silica capillaries (100 µm inner/360 µm outer diameter) with a laser puller (P-2000, Sutter Instruments) and packed with either 5 µm or 3 µm C_18_ particles (Reprosil-Pur 120 C_18_ AQ, both Dr. Maisch).

For LTQ Orbitrap Velos analyses, peptides were loaded onto the analytical column (20 cm length, 5 µm particles) with 99% buffer A (5% DMSO, 95 % H2O, 0.1% FA) 1% buffer B (95% ACN, 5% DMSO and 0.1% FA) at 1 µl/min for 20 min and eluted with a 30-min linear gradient from 1-35% buffer B at a flow rate of 400 nl/min. Eluting peptides were ionized in the positive ion mode and MS1 scans were recorded in the Orbitrap (OT) analyzer from m/z 300-2000 at a resolution of 60,000. The ten most abundant MS1 precursor ions were selected for collision-induced dissociation (CID) fragmentation with a normalized collision energy (NCE) of 35, MS2 spectra were acquired in the linear ion trap, and dynamic exclusion was set to 30 s. For Orbitrap Fusion Lumos analyses, samples were either loaded directly to the analytical column (35 cm, 3 µm particle size) with 99% buffer A (100% H2O, 0.1% FA) 1% buffer B (90% ACN, 10% H2O, 0.1% FA) at a flow rate of 600 nl/min for 25 min, or onto a trapping column (Acclaim PepMap C_18_ 5 µm 100 Å, 160454, Thermo Fisher Scientific) with 0.5% TFA at a flow rate of 10 µl/min for 8 min. Peptides were eluted with linear gradients of 30-, 60- or 120 min from 1-35% buffer B at 300 nl/min and ionized in the positive ion mode. For DDA analyses, MS1 scans were acquired from m/z 250-2000 at a resolution of 60,000 in the OT analyzer with an AGC target setting of 4×10^5^ and a maximum injection time set to 50 ms or to “automatic. The most abundant MS1 precursor ions (+2 to +7) were isolated in the quadrupole (m/z 1.2-1.6), fragmented by higher energy collisional dissociation (HCD) with an NCE of 30 in the top speed mode (2-3 s cycle time), and excluded for 15- 60 s from fragmentation. MS2 fragment ion spectra were acquired at a resolution of either 15,000 or 30,000 in the OT mass analyzer using a AGC target setting of 5×10^4^ with maximum injection times between 22-100 ms. For PRM experiments, MS1 scans were recorded from m/z 200–2000 and the maximum injection time set to 118 ms. Up to 140 ms maximum injection time was used in the PRM experiment, recording MS2 spectra at a resolution of 60,000.

For MRM experiments conducted with the QTrap5500 mass spectrometer, peptides were injected at a flow rate of 10 µl/min in 99.4% buffer A (98.9% H2O, 1% ACN, 0.1% FA) 0.6% buffer B buffer B (89.9% ACN, 10% H2O, 0.1% FA) for 7 min onto a trapping column (Symmetry C_18_ 5 µm 100 Å, 186007496, Waters), followed by their elution onto, and separation by, an analytical column (M-Class Peptide BEH C_18_ 1.7 μm,130 Å, 186007484, Waters). Peptides were eluted with a linear gradient of 3-40% buffer B for 177 min at a flow rate of 200 nl/min and a column temperature of 35 °C. For the QTrap 6500+ mass spectrometer, samples were loaded at a flow rate of 20 µl/min with 0.1% TFA for 4 min onto a trapping column (Acclaim PepMap C_18_ 5 µm 100 Å, 160454, Thermo Fisher Scientific) followed by their elution to, and separation by, the analytical column (35 cm, 3 µm particle size) at 300 nl/min with a linear gradient from 7% buffer B (90% ACN, 10% H2O, 0.1% FA) 93% buffer A (100% H2O, 0.1% FA) to 35% buffer B in 30 or 60 min. Both QTrap instruments were operated in the MRM mode (low mass region) using unit resolution for the quadrupole analyzers. Peptides were ionized in the positive mode. MRM-initiated detection and sequencing (MIDAS) experiments were performed in the unscheduled mode, with 20 ms dwell time per transition and a maximum of 60-75 transitions per sample injection [75]. Collision energies (CEs) were optimized for a selected set of peptides (individual transitions, step size 2.5 V) using Skyline (21.1.0.146, skyline.ms [76, 77]), the default CE parameters and optimized values were incorporated to the assay [78]. Scheduled MRM experiments were conducted with 3 s cycle time and detection windows of 3 min defined.

### MS data processing and analysis

Acquired DDA files were processed with Proteome Discoverer (2.5.0.400, Thermo Fisher Scientific) utilizing Mascot as search engine (v2.6.1, Matrix Sciences) or Mascot as standalone application. Searches were performed against a custom QconCAT database (Table S1), or a combination of the QconCAT database and SwissProt E. coli (4403 entries, release 07/2023), along with a database containing common contaminants [79], with a concatenated decoy generation strategy. For database searches, fully tryptic peptides with up to one (labeling efficiency study and library generation) or three (missed cleavage study) missed cleavage sites were considered. The mass tolerance was set to 10 ppm (MS1) and 20 ppm (MS2) for data acquired on the Orbitrap Fusion Lumos and 10 ppm (MS1) and

0.6 Da (MS2) for data acquired on the LTQ Orbitrap Velos. MRM data were converted to mgf files using ProteoWizard (3.0.221466 [80]) and searches performed with a 1.2 Da MS1 and 0.6 Da MS2 mass tolerance. For all searches, propionamide (C) or carbamidomethyl (C) were defined as static modifications and oxidation (M) as well as acetyl (protein N-term) as variable modifications. Heavy amino acid labeling with ^13^C_615_N_2_ (K) and ^13^C_615_N_4_ (R) was set as fixed modification for the analysis of biological samples and as variable modification for labeling efficiency analyses. Identified peptide spectrum matches (PSMs) were validated based on q-values with Percolator [81] using a 1%/5% (strict/relaxed) target false discovery rate (FDR). Combined PSMs were aggregated to peptides and proteins applying the principle of strict parsimony, and a 1% FDR filter was applied at the peptide and protein level.

MRM data were analyzed with Skyline. Spectral libraries were built with the BiblioSpec plugin of Skyline applying a cut-off score of 0.95 (settings: filter for documented peptides, exclude ambiguous matches). When both QTrap and Orbitrap MS2 spectra were available for one peptide, QTrap data were used. For peptides with no available spectra, MS2 spectra were generated in silico using PROSIT [82] (NCE27, default settings with intensity prediction model: publication_v1). MRM data quality was monitored by manual inspection and by using the AutoQC loader application (22.1.0.122, [83]) in combination with Skyline. Sample-specific models were generated with mProphet [84] and trained using the 2nd best peak option, excluding negatively contributing features. Inconclusive stable isotope-labeled peptide standard traces were deactivated and excluded from the following data analysis. In the case of interfering or low/noisy endogenous sample peptide traces, the theoretically highest fragment ion trace was used, excluding other transitions („Not quantitative“ feature). Skyline peptide quantity exports were processed in MS Excel and R (v4.2.2 [85]) using the reshape2 (v1.4.4), dplyr (v1.1.3), tidyr (v1.3.0), and stringr (v1.5.0) packages. For peptides with both oxidized and non-oxidized variants, the summed values of both versions were used. Signal-to-noise (S/N) ratios were calculated as ratio of total fragment/total background areas. Peptide quantifications were classified into three categories as follows: “A” for S/N ratios ≥ 3 of both endogenous and stable isotope-labeled peptide, “B” for S/N ≥ 3 of stable isotope-labeled peptide, and S/N < 3 for endogenous peptide, and “C” if stable isotope-labeled peptide had S/N < 3. C peptides were not considered for protein quantification, employing two modes for removal: medium and stringent. Medium removal excluded class C peptide quantifications in the affected sample/condition, requiring at least two remaining quantifications per sample/condition. Stringent class C peptide removal eliminates the peptide quantification entirely from the data matrix across all samples/conditions. Unless otherwise stated, median absolute protein amounts are reported for all replicates per sample and/or condition and the variance was assessed via the robust standard deviation (rSD), calculated as 1.4826 x median absolute deviation (MAD) which is defined as: (çX_i_ - median_j_(X_j_)ç). The robust coefficient of variation (rCV) as relative measure of variance was calculated as: rSD/median [55]. For visualization the R packages ggplot2 (v3.4.3), gridExtra (v2.3), RColorBrewer (v1.1.3) and ComplexHeatmap (v3.2), as well as GraphPad Prism (10.1.2) were used.

### Immunofluorescence analysis

Cells were seeded on 15 mm coverslips in 12-well plates at a density of 1.5×10^4^ and grown for 24 hrs. For Ctsd visualization, cells were incubated with SiRLysosome (1 µM, SC02, Tebu-bio) for 2 hrs prior to fixation. Cells were fixed in 4% paraformaldehyde (Sigma Aldrich), 1x PBS for 25 min at 37 °C, washed 3x with 1x PBS, permeabilized with 0.1% Saponin (S7900, Sigma Aldrich/Merck) 2% BSA in 1x PBS at RT for 10 min, and subsequently incubated with anti LAMP2 (1:400, clone ABL-93, DSHB) in blocking solution (0.1% Saponin, 2% BSA in 1x PBS) at 4 °C overnight. The following day, cells were washed (1x PBS) and incubated in blocking solution with fluorophore-conjugated secondary antibody (1:500, Cy3 AffiniPure Goat Anti-Rat, 112-166-062, Dianova) at RT for 1 hr. Washed cells were mounted with 1x ROTI Mount FluorCareDAPI (HP20.1, Carl Roth) to specimen slides. Fluorescence microscopy imaging was performed with an Axio Observer 7 microscope in combination with a 63x objective (Axiocam 705 mono, both ZEISS). Images were deconvolved, processed with maximum intensity projection (v3.4, Zen Bluepro, ZEISS), and lysosomes were counted with segmentation masks generated using Ilastik [86]. A subset of images was manually annotated based on Lamp2 signals to train a custom classification model for lysosomes, the classifier was applied to the full dataset, and the Fiji package [87] was used for the definition of lysosome masks and lysosome counting. For mask definition, particle edge smoothing, hollow particle filling, and watershed segmentation was applied. The Mander’s coefficient was calculated using the Fiji JaCoP plugin.

### Quantitative RNA analysis of primary cell types

RNA was isolated from cultured lung fibroblasts, macrophages, and osteoblasts (Monarch Total RNA Miniprep Kit, NEB) and 1 μg of total RNA from pooled samples was reverse transcribed (High-Capacity cDNA Reverse Transcription Kit, Thermo Fisher Scientific). For quantitative PCR (qPCR), TaqMan gene expression assays with pre-designed probes and primers for lysosomal enzyme cDNAs were used [88, 89]. All reactions were run in technical triplicates using the primaQUANT qPCR Master Mix (Steinbrenner) on a QuantStudio 1 PCR system (Thermo Fisher Scientific). Relative mRNA levels were normalized to Gapdh within the same cDNA sample, using the 2–ΔΔCT comparative method.

### Protein copy number determination and protein distribution analysis

The degree of lysosomal localization for individual proteins was calculated as follows:

Determination of protein copies per lysosome in whole cell samples:

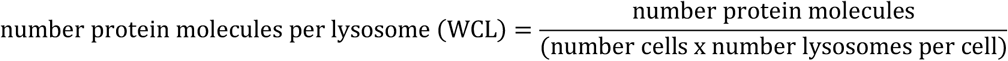

with: number of cells determined from cell counting and protein assays and number of lysosomes determined from immunofluorescence analysis.

1. B) Determination of protein copies per lysosome in lysosome enriched fractions:
2. The number of lysosomes in the lysosome enriched fraction was determined as:

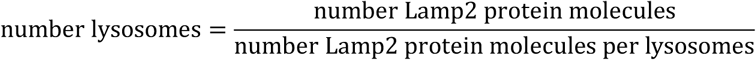

with: number Lamp2 molecules per lysosome: 1275 (determined from immunofluorescence analysis).

1. 2. The number of protein molecules present at one lysosome was determined as:

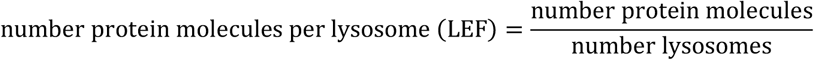

with: number of lysosomes as determined in (1).

1. C) Determination of protein molecule distribution between lysosomal and non-lysosomal localizations.
2. The number of non-lysosomal located protein molecules was determined as:

number protein molecules per lysosome (WCL) − number protein molecules per lysosome (LEF)

1. 2. The relative proportional distribution (%) for lysosomal-located (as determined in B2) and non-lysosomal located protein molecules (as determined in C1) was calculated as proportion of the number protein molecules per lysosome (WCL) determined in A.

## RESULTS AND DISCUSSION

In order to enable absolute quantification of the lysosomal proteome across different types of mouse samples, we developed a strategy based on the generation of stable isotope-labeled standard peptides and a targeted MRM-MS assay. We included proteins which were previously reported to be located either in the lysosomal lumen or its membrane, and hence likely to be important for lysosomal function.

### Generation of Absolutely Quantified Protein Standards

Based on a manually curated list of known lysosomal proteins from our group [9] and proteins which were identified in unbiased screenings to be located at lysosomes (Table S1), we chose 155 proteins. For those, we selected tryptic surrogate peptides from both in house and published datasets [56–59], identifying > 6300 candidates. Based on their biochemical and physiochemical criteria relevant for standard peptide release from QconCATs and MS detection, we manually selected 414 peptides covering 144 proteins (Fig. 1A, Table S1). Subsequently, we grouped them based on their average abundance in large scale proteomics studies [56–59] and distributed them to 12 QconCAT proteins. We then optimized the peptide order to yield full tryptic digestion of individual QconCATs, added a C- terminal minimally permutated peptide analog (MIPA, [61]) as internal standard to enable absolute quantification (Table S1), and an N-terminal His-tag for affinity purification. Following reverse translation of the final amino acid sequence and optimization for bacterial expression, synthetic constructs were generated by gene synthesis and cloned into bacterial expression vectors (Supplementary file 1). Finally, we expressed individual QconCATs in minimal media supplemented with stable isotope-labeled amino acids, purified them, and confirmed protein expression by SDS-PAGE and western blotting (Fig. S1).

**Figure 1:**
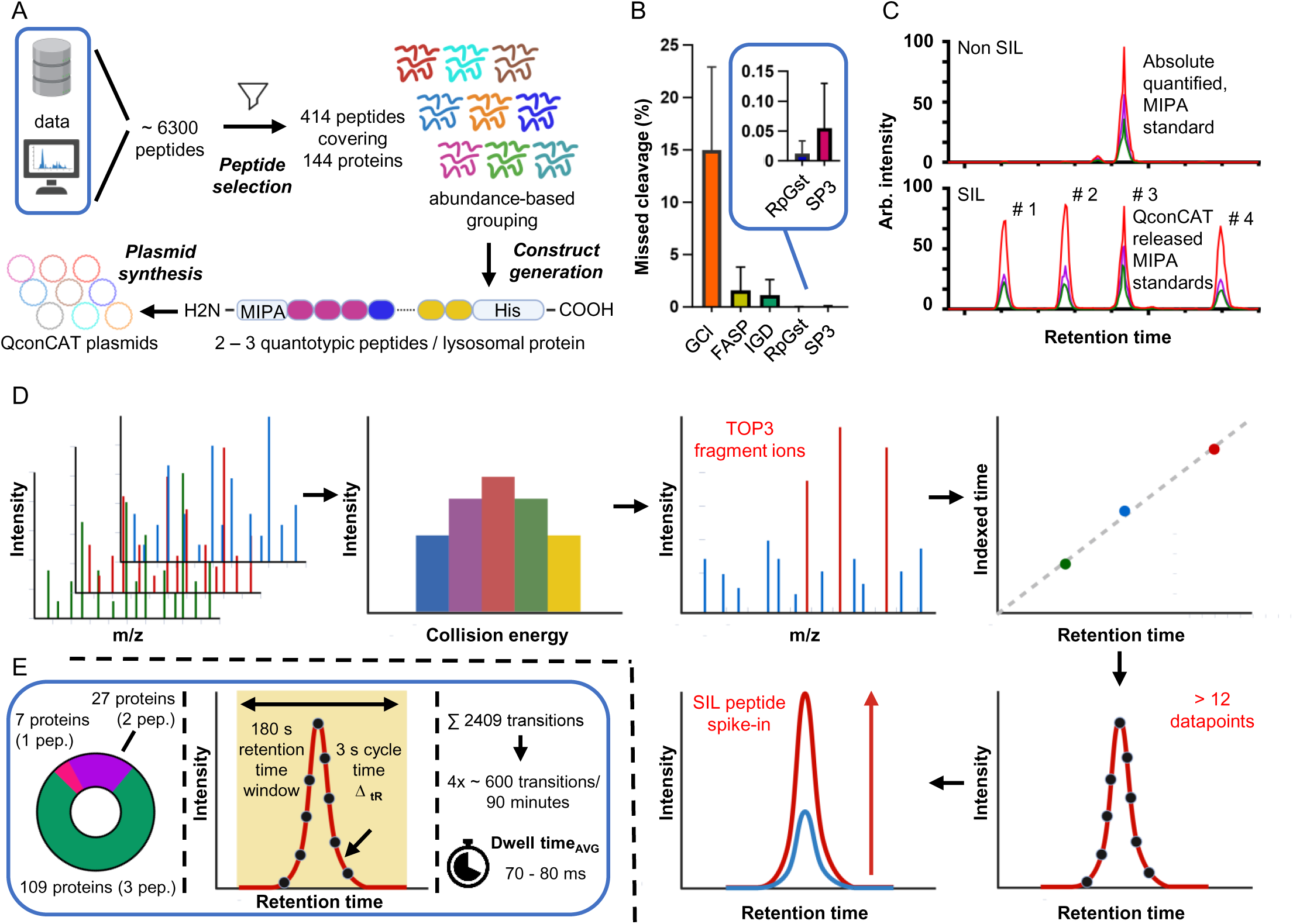
Design, evaluation and re-quantification of QconCAT protein standards. A) Schematic overview of the QconCAT workflow designing artificial QconCAT fusion proteins. Key steps include the selection and assembly of proteotypic peptides into a protein construct optimized for efficient proteolytic digestion. Following construct optimization and modification, synthetic genes are generated. B) Comparison of strategy-specific digestion performance. Reported are the cumulative missed cleavage rates (%) averaged for all 12 QconCAT protein standards per digestion strategy tested. C) Schematic representation of a multiplexed, absolute re-quantification experiment allowing re-quantification of four digested SIL QconCAT proteins with one non SIL synthetic MIPA peptide. **MRM assay development and optimization.** D) Empirical data from MS experiments gathered are compiled into reference libraries (MS2 spectrum and normalized retention time). Targeted MS assay development comprises the determination and optimization of peptide, as well as assay-specific parameters in a multistep process. E) Key metrics of final MRM assay. Shown is the protein coverage by the number of peptides as well as assay-specific parameters used for data acquisition. Abbrev.: CE: collision energy. SIL: stable isotope labeled. Figure 1 A and D were generated using BioRender.

To utilize QconCAT-derived standard peptides for absolute quantification experiments requires the complete and efficient release of the individual reference peptides contained in the protein standard. We therefore evaluated the performance of different digestion strategies including bead-based single-pot, solid-phase-enhanced sample-preparation (SP3) [65], in solution digestion using guanidinium chloride [69] or RapiGest [66, 70], Filter Aided Sample Preparation (FASP) [68], and in gel digestion (IGD) [67]. Based on untargeted LC-MS/MS analyses of tryptic digests of all 12 QconCATs, we determined the average missed cleavage rate, identifying lowest values with SP3, RapiGest, and IGD, and an average labeling efficiency of > 98% (Fig. 1B, Fig. S2, Table S1). As IGD enables further to select QconCATs based on the molecular weight of the protein, eliminating truncated protein standards that result in false abundances due to the lack of distinct peptides, we chose this strategy for the generation of stable isotope-labeled lysosomal peptide standards (SILLPS). Due to the compatibility with large sample amounts, we selected RapiGest digestion for the proteolytic digestion of biological samples.

### MRM Assay Development

To determine the quantity of individual QconCAT-derived SILLPS, we obtained absolutely quantified synthetic MIPA reference peptides, which we used as internal standards for re-quantification in MRM analyses (Fig. 1C, Table S1, Fig. S3). Based on these data, we prepared master mixes containing equimolar amounts of all QconCATs, optimized collision energies for peptide fragmentation, defined reference MS2 spectra for spectral library generation using DDA, PRM, and MIDAS [75] runs, and selected the three most abundant, interference-free fragment ions for the final MRM assay. We then determined peptide-specific, normalized retention times using the indexed RT (iRT) concept [90]. To determine SILLPS spike-in and on-column sample loading amounts, we utilized mixtures of cell line digests (fibroblasts, osteoblasts, osteoclasts, and macrophages). We aimed at a range of up to two orders of magnitude between SIPPLS and internal peptides, to enable accurate quantification with the utilized QTrap6500+ mass spectrometer. This resulted in two different spike-in doses for individual QconCATs (Fig. 1D). Finally, we determined optimal MS parameters, MRM assay size, gradient lengths, and numbers of runs to cover the full assay. Peptides which did not meet our criteria for robust quantification in these analyses were removed from the assay.

The final MRM assay targets 388 lysosomal protein-derived peptides and their respective SILLPS covering > 95% of the included 143 lysosomal proteins with two or three surrogate peptides through ∼2400 individual transitions (Fig. 1E, Table S1). To enable acquisition of a sufficient number of data points across individual chromatographic peaks, we split the assay into four separate LC-MRM-MS analyses, and defined the retention time windows as well as cycle and dwell times accordingly.

### Lysosomal Protein Copy Number Determination in Mouse Embryonic Fibroblasts

Several factors are crucial for correct lysosomal function. In order to obtain functional and stable organelles, several types of membrane proteins are needed. Structural proteins with heavily glycosylated luminal domains, such as Lamp1/2 [91] or Glmp [92], stabilize the lysosomal membrane [93] and prevent lysosomes from self-digestion [94]. The vacuolar ATPase (vATPase [95]), a 1 MDa multi protein complex, pumps protons across the lysosomal membrane, generating its acidic pH. For the export of degradation products to the cytosol and the bidirectional transport of ions such as Ca^2+^, K^+^ or Cl^-^ a variety of membrane-embedded transporters, channels, and exchangers exist [96–98]. The degradative function of the lysosome, on the other hand, is facilitated by a multitude of luminal hydrolases, which can be broadly classified based on their substrates (e.g. proteases, lipases, nucleases, etc.) [99].

In order to investigate the abundance of these individual protein classes, we initially used our assay to determine lysosomal protein copy numbers in whole proteomes of mouse embryonic fibroblasts (MEFs), one of the most commonly used immortalized mouse cell lines (Fig. 2A). To determine protein copy numbers per cell, we quantified the average protein content of MEF cells by a combination of cell counting and protein assays to be 0.22 ng/cell (Fig. S4), and combined this information with the SILLPS spike-in-derived values (Table S2). In this copy number/cell information for 141 MEF lysosomal proteins, we observed a dynamic range of more than three orders of magnitude between the lowest and highest abundant lysosomal proteins, respectively (Fig. 2B). The two lysosomal membrane proteins Lamp1/Lamp2, the small cholesterol binding protein Npc2, and the putative lysosomal protein Pituitary tumor-transforming gene 1 protein-interacting protein (Pttg1ip [100]) presented the highest lysosomal proteins expressed in MEF, and the only proteins in our dataset exceeding 500,000 copies/cell. These findings align closely with other studies that identified these proteins to be highly abundant [37], and based on these determinations Lamp1 and Lamp2 contribute together ∼19% of lysosomal membrane protein molecules (based on copy numbers), equivalent to ∼12% of lysosomal membrane protein biomass and 7% of total lysosomal biomass. The four lowest expressing proteins comprised the three hydrolases Aoah, Ctss, and Neu4, as well as the oligopeptide transporter Abcb9.

**Figure 2:**
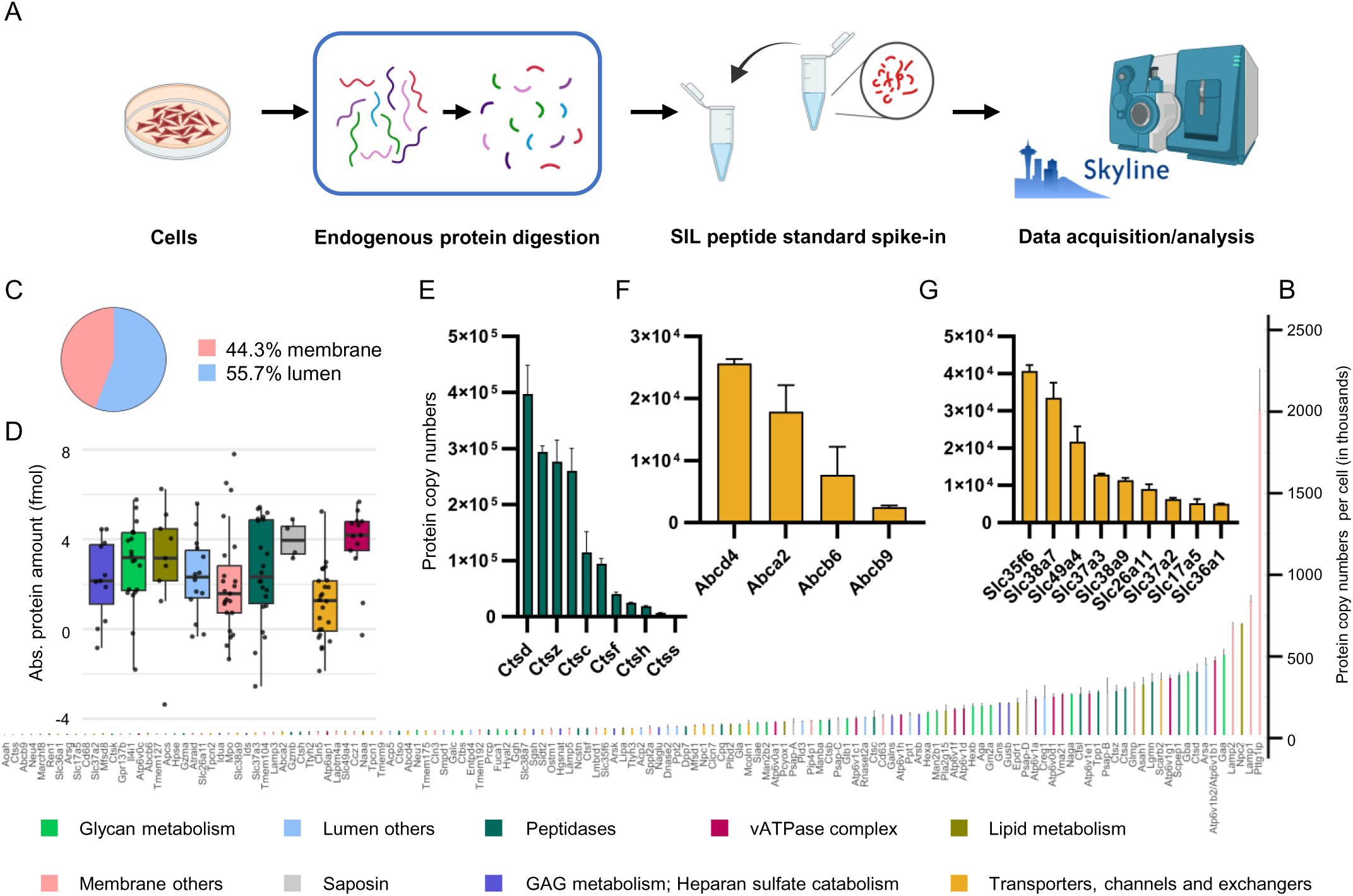
Absolute quantitative analysis of lysosomal proteins in mouse embryonic fibroblasts. A) Workflow for absolute quantification experiments. Harvested cells are proteolytically digested and defined peptide amounts are mixed with known amounts of SIL QconCAT peptide standard. Endogenous peptide/protein amounts are derived from absolute quantified peptide standard amounts. B) Determined protein copy numbers for 141 proteins per cell. Proteins are associated with nine different lysosomal classes/categories. C) Distribution of cumulative protein copy numbers. D) Distribution of log2-transformed, absolute protein levels associated with different lysosomal classes/categories. Boxes display the interquartile range (IQR), with each box indicating the median protein expression level for the classes/categories. Whiskers extend to the smallest and largest values within 1.5 times the IQR from the lower and upper quartiles. E, F and G) Copy numbers of lysosomal protein molecules per one MEF cell. Median values across biological replicates (n=3) are reported; error bars indicate the robust standard deviation (rSD). Abbrev.: MEF: mouse embryonic fibroblast.

Considering cumulative protein copy numbers of all lysosomal proteins, we observed a similar distribution in summed abundance of luminal (55.7%) and membrane (44.3%) proteins, respectively (Fig. 2C). The proteins forming the vATPase complex showed the highest median abundance (∼30% of lysosomal membrane proteins) and lowest spread, which is not surprising considering its crucial role for lysosomal function and defined stoichiometry [101], as well as saposin A/B/C/D (Fig. 2D), which originate from a common precursor protein and are released by proteolytic cleavage [102]. With respect to lysosomal hydrolases, the four biggest functional groups account for > 70% of lysosomal luminal protein amount, namely 29% glycosidases (23 proteins), 28% proteases (22 proteins), 10% sulfatases (8 proteins), and 6% lipases (5 proteins). The spread of their absolute protein abundance core distribution, represented by the interquartile range (25th to 75th percentile) for glycosidases, sulfatases, and lipases behaved verily similar, while peptidases markedly exceeded the other groups (discrepancy of > 250-fold between highest/lowest members, Fig. 2D). For the latter, we further investigated the sub- group of cathepsins (11 proteins, Fig. 2E) which were distributed across the complete dynamic range of peptidase expression levels. For most protein classes associated with the lysosomal membrane, we observed a similar spread, but marked variations in their median expression, with lowest values for transporters, channels, and exchangers (12% of cumulative lysosomal membrane proteins with a class-specific dynamic range of ∼140-fold). Within distinct sub-groups of this class, however, we observed significantly lower dynamic ranges than for hydrolases, such as 8-fold for ATP-Binding Cassette (ABC) transporters (4 members, Fig. 2F), and 10-fold for solute carrier (SLC) transporters (9 members, Fig. 2G), respectively.

### Estimation of lysosome-specific protein copy numbers

While data obtained from whole cell lysates (WCL) allow for the estimation of total protein expression levels per cell, they fail to provide accurate information if this amount of protein is indeed present at lysosomes. While certain lysosomal luminal proteins are known to be solely lysosomal, for several lysosomal (membrane) proteins, such as SLC transporters, a dual localization was shown [103]. To enable estimation of the subcellular distribution of lysosomal proteins, we enriched lysosomes using superparamagnetic iron oxide nanoparticles (SPIONs, Fig. 3A) [45] and analyzed the resulting lysosome-enriched fractions (LEFs) with our MRM assay in the presence of SILLPS (Table S2). As it is difficult to determine the exact number of lysosomes contained in LEFs, and hence to calculate protein copy numbers per organelle analogous to WCL data, we devised an alternative strategy based on a combination of fluorescence microscopy and protein normalization. First, by co-staining analyses for Lamp2 (antibody-based) and hydrolytically active Ctsd (SirLysosome [104]), we determined the number of hydrolytically active terminal lysosomes per MEF cell to be 482 under our cell culture conditions (95% of Lamp2-positive structures contained active Ctsd, Fig. 3B). At the same time, we did not observe any non-lysosomal Lamp2 intensity, placing all molecules at this organelle, and, based on our SILLPS, locating ∼1300 Lamp2 molecules to each lysosome (in case of uniform distribution). We utilized this value to determine that one LEF from two 10 cm plates of MEFs contains ∼7.6×10^7^ lysosomes, and to calculate individual protein copy numbers per lysosome both in WCLs and LEFs (Table S2). For LEFs, we observed a similar dynamic range as for the WCL with marked differences for individual proteins (Fig. S5). The most abundant molecules in LEFs were saposins B and D, Lamp2, Cd63, and Pttg1ip, matching the pattern for Lamp2 and Pttg1ip observed in whole cell lysates, but a marked increase in relative abundance for the saposins and Cd63. Interestingly, also in these analyses the putative lysosomal protein Pttg1ip, for which no evidence for a lysosomal function exists to date, was among the five most abundant proteins.

**Figure 3:**
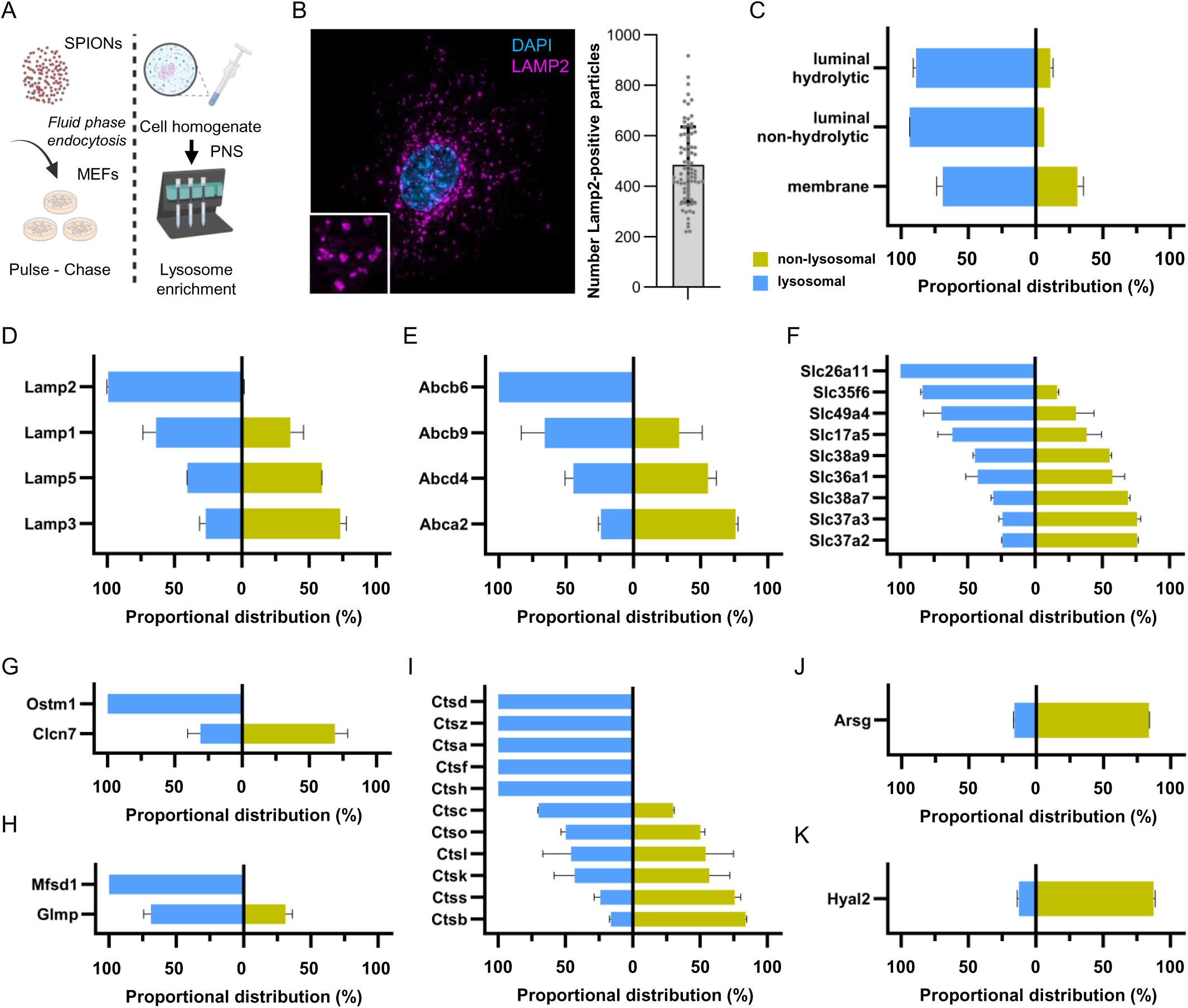
Lysosomal protein distribution analysis. A) Schematic workflow of intact lysosome isolation using paramagnetic beads. Lysosomes loaded with magnetic particles are retained and isolated from cell homogenate. B) Lysosome-associated membrane protein 2 (LAMP2) immunofluorescence staining in MEF cells (left). Counting of LAMP2-positive markers (n=79, right). C-K) Proportional distribution of lysosomal protein molecules normalized to one cell. The distribution of protein molecules localized in/at the lysosome (lysosomal, left) and outside the lysosome (non- lysosomal, right) is shown. The sum of individual protein molecule numbers per category was used in C). Proportional distributions are shown for D) lysosome-associated membrane proteins, E) ATP- binding cassette transporters, F) solute carrier proteins, G) lysosomal dipeptide transporter (Mfsd1) and glycosylated lysosomal membrane protein (Glmp), H) osteopetrosis-associated transmembrane protein 1 (Ostm1) and H(+)/Cl(-) exchange transporter 7 (Clcn7), I) cathepsin proteins, J) arylsulfatase G (Arsg), and K) hyaluronidase-2 (Hyal2). Figure 3A was generated using BioRender.

### Protein distribution analysis identifies differential localization for individual lysosomal proteins

Based on the correlation of lysosome-normalized protein copy numbers for WCL and LEF samples, we investigated the spatial distribution of the individual proteins covered by our assay, revealing that ∼85% of total lysosomal protein molecules also localized to this organelle (Fig. S6A). We then addressed individual lysosomal protein classes based on their localization, identifying sub-group specific distribution patterns, with higher fractions of non-lysosomal localization for proteins which are located at the lysosomal membrane (Fig. 3C). These findings indicate that the majority of lysosomal luminal proteins are directly transported to lysosomes, presenting their final destination, while membrane proteins are either retained to a certain degree in the ER and/or Golgi, or permanently localize to entirely different subcellular compartments. For individual proteins, we observed that 79 of them showed a primarily lysosomal localization, while for the remaining 62 proteins more than one-third were present at another subcellular location (Fig. S6B).

We observed, for example, full lysosomal localization of Lamp2, while Lamp1, 3, and 5 only localized with 64, 27, and 41% (Fig. 3D), which is in line with previous reports placing only a sub-fraction of Lamp1 at degradative lysosomes [105, 106]. The vATPase complex presented with a quite heterogenous pattern for the presence of individual subunits with, surprisingly, the highest and lowest rate for two members of the V1 subcomplex (Fig. S6C). As the complex exists at lysosomes in a defined stoichiometry, it is possible that such subunits with a high percentage of lysosomal localization present the rate-limiting members for complex formation, while such with a high non-lysosomal fraction are either present in excess or fulfil additional functions in the cell.

### Lysosomal localization of membrane proteins

Broadly, lysosomal membrane proteins can be classified in such without any role for the exchange of metabolites, as well as transporters, channels and exchangers. Of the second class, five proteins showed exclusive lysosomal localization (Figure S6D), from which only one protein each belonged to the two biggest groups, the ABC and SLC transporters (Fig. 3 E, F). The sodium-independent anion exchanger Slc26a11 and the porphyrin transporter Abcb6 localized in our data solely to the lysosome. While both of these proteins have been shown in other cell lines to also localize to other subcellular compartments, in our MEF data they show exclusive presence at the lysosome. This could be related to the unique properties of MEF lysosomes, which we showed previously to present with a composition differing from other commonly used cell lines [37]. With respect to SLC transporters, interestingly, the proteins with the smallest percentage of lysosomal localization (∼25%) were the two members of the SLC37 sugar-phosphate exchangers, which were initially considered to be ER-resident proteins [107, 108]. Those SLC transporters which are involved in amino acid transport (Slc38a9 [109], Slc36a1 [110], and Slc38a7 [111]) which have been described as integral components of the lysosomal amino acid-sensing machinery and play a critical role in amino acid sensing by mTORC1 [109, 110], showed a higher degree of lysosomal localization. We still found, however, > 50% of each of these transporters to be present in non-lysosomal regions of the cell.

The lysosomal chloride-proton exchanger Clcn7 and its beta subunit Ostm1 present with unique properties in this context. It was shown previously, that Clcn7 is present in an ER-resident pool and is only transported to lysosomes upon assembly with Ostm1 [112]. In accordance with these findings, we detected 100% of Ostm1 to be located at lysosomes, while only 31% of Clcn7 was present at the organelle (Fig. 3G). A similar picture presented for Mfsd1 and its accessory subunit Glmp, which were recently demonstrated to be act as general dipeptide uniporter in lysosomes [92, 113]. While all copies of Mfsd1 localized to lysosomes, only ∼70% of Glmp were present there (Fig. 3H), which could be related to a similar relationship.

### Lysosomal localization of luminal proteins

The large majority of lysosomal luminal proteins are hydrolases, and their confinement to this subcellular compartment is an essential mechanism to restrict their activity to a secure environment, as mis-localization would result in self-digestion of the cell [114]. It is, therefore, not surprising that we detected a significantly higher degree of lysosomal localization for this class of proteins relative to such located at the membrane (Fig. 3C), and that the majority of fully lysosome-localized proteins are located in the lumen (Fig. S6B). Interestingly, with respect to individual classes of hydrolases, such related to glycan metabolism showed the highest degree of lysosomal localization, followed by peptidases, proteins involved in lipid degradation, and those facilitating the degradation of glycosaminoglycans/heparan sulfate (Fig. S6E, F, G. H). This could be related to non-lysosomal functions of certain proteins or their mode of activation, as it is conceivable that the ∼50 lysosomal hydrolases which are not activated by proteolytic cleavage, are more strictly restricted to this compartment.

Peptidases present a distinct group in this context, as all of them catalyze the cleavage of catalytic bonds in a rather unspecific way [115], and hence present with a certain redundancy [39], while other lysosomal hydrolases engage in highly specific degradative reactions. Four of the five highest abundant peptidases (Ctsd, Lgmn, Ctsa, and Ctsz) show exclusive lysosomal localization in our data and the fifth protein of this group, Scpep1 is located with ∼80% at the organelle (Fig. 2B, Fig. S6F), placing the majority of peptide hydrolytic activity in this compartment (more than 60% of total peptidase copy numbers). We observed the opposite behavior for Ctsk, Ctss, and Ctsb of which only 43%, 24% and 16% localize to the lysosome (Fig. 3I). Notably, especially Ctsb was shown in numerous studies to exert non-lysosomal functions with involvement in tissue invasion, apoptosis [116], or inflammation [117], which is well in line with a pool of this protein that is localized at other subcellular compartments.

Two other hydrolases which we detected to be mainly of non-lysosomal localization are Arsg (17% lysosomal, Fig. 3J) and Hyal2 (19% lysosomal, Fig. 3K), which are involved in catabolism of the large glycosaminoglycans heparan sulfate and hyaluronan, respectively. Arsg is responsible for the degradation of O-sulfated N-sulfoglucosamine residues in heparan sulfate and was demonstrated to localize to pre-lysosomal compartments associating closely with organelle membranes, likely those of the endoplasmic reticulum (ER) [118]. Contradicting reports exist regarding the proteolytic processing of ARSG, with both unprocessed and processed protein forms being associated with hydrolytic activity [118, 119]. For Hyal2, on the other hand, there is an ongoing debate regarding its localization [120–123], which our data clearly identifies as mainly non-lysosomal in MEFs. The fact, that both of these hydrolases, which are involved in the degradation of constituents of the extracellular matrix, rank among the least lysosomal-localized proteins, and that they may not require proteolytic activation presenting with extralysosomal activity [118, 119], could indicate a function at another subcellular compartment, possible in the maturation of molecules before their transport to the extracellular space.

### Cell type specific lysosomal protein expression patterns show group-specific similarities

As MEF cells, like all other commonly used immortalized cell lines, have been grown in tissue culture for countless passages, it is questionable in how far their proteome is a realistic representation of cells growing in solid tissues; as it was shown that even within the same cell type grown at different laboratories major proteome differences exist [124]. Primary cells present a more realistic picture and often the only option to obtain a sufficiently pure population of the cells of interest, as the enrichment of distinct cell types from tissue by Fluorescence-Activated Cell Sorting (FACS) or Magnetic-Activated Cell Sorting (MACS) is often problematic. They are, however, often not available in the quantities needed to enrich lysosomes and establishment of lysosome enrichment conditions by Lyso-IP or SPIONs is laborious and requires extensive growth in culture, which may again alter their cellular proteome, e.g. through the overexpression of tagged membrane proteins [125]. Our MRM assay presents an ideal solution to investigate such cells and we applied it to primary fibroblasts (as direct comparison to MEFs) as well as three cell types with distinct lysosomal features, due to varying roles of lysosomes in them. In detail, we investigated lung fibroblasts (LFs), osteoblasts (OBs), osteoclasts (OCs), and macrophages (MPs), which were either cultured directly from tissue homogenates or generated through differentiation of pluripotent hematopoietic stem cells [126–128].

After determination of lysosomal protein copy numbers in all cell lines using LC-MRM-MS in combination with SILLPS, we initially calculated the amount of lysosomal proteins present in each cell line (Table S3) and compared them to the data generated from MEFs. For MEFs, we observed a markedly lower cumulative amount of lysosomal proteins relative to primary cells, with differences of up to 4-fold, and highest abundances of lysosomal proteins for OBs, OCs, and MPs, which is in line with their high lysosomal activity (Fig. 4A). To investigate if this change is due to a specific portion of the lysosomal proteome, we compared protein intensity distributions of lysosomal luminal and membrane proteins for each cell line (Fig. 4B), revealing a higher relative contribution of hydrolases in cells with higher lysosomal activity (∼80% luminal and 20% membrane proteins, (Fig S7A). To further address differences in individual lysosomal proteins, we performed binary comparisons by Pearson correlation, revealing high levels of similarity for the majority of lysosomal proteins of MEFs and LFs, as well as of, OBs, OCs, and MPs, with pronounced differences between the two groups of cells (Fig. S7B). Also, unsupervised hierarchical clustering showed similar patterns within these groups and, interestingly, failed to distinguish between MPs and OCs, indicating a high similarity of their lysosomal proteome. (Fig. 4C). To further investigate the differences between the individual cell lines, we compared the summed abundance of individual classes of proteins (Fig. 4D, Fig. S7C). This revealed that especially the amount of vATPase complex members, peptidases, and saposins were markedly higher in OBs, OCs, and MPs which is indicative of a higher degradative capacity, and possibly substrate turnover, of these cells. Also, these data showed that the ratio of the vATPase to other structural membrane proteins is conserved across primary cell lines, implying a similar organization of the lysosomal membrane in these cells, despite their varying luminal content.

**Figure 4:**
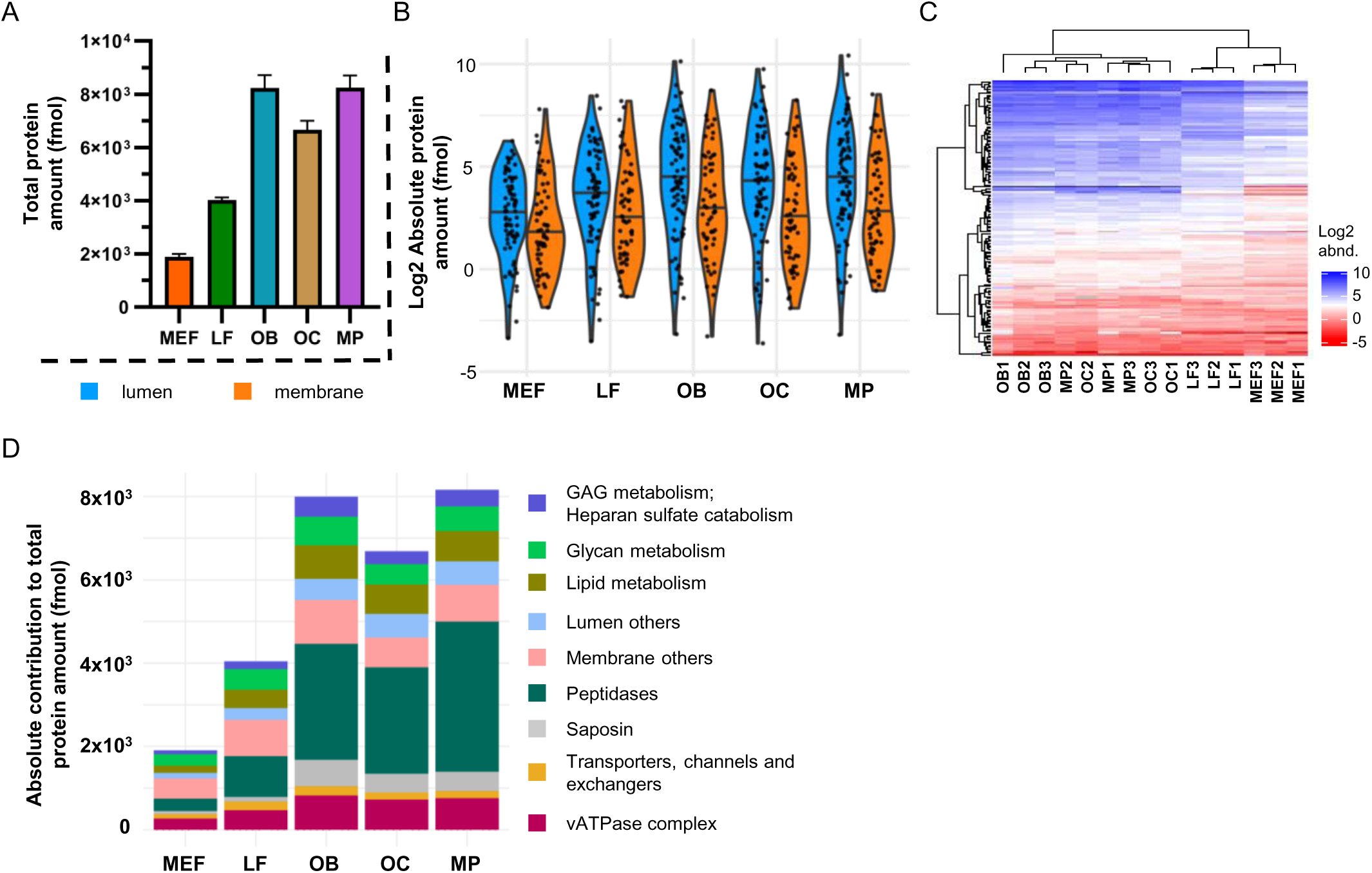
Cell type-specific analysis of lysosomal protein expression. A) Comparison of summed protein abundances across cell types. B) Intensity distribution of luminal and membrane protein abundances. C) Heatmap of hierarchical clustering (complete linkage, Euclidean distance, column- and row-wise clustering) for 141 proteins. Missing values are indicated in grey. D) Absolute contribution of protein classes to the total lysosomal protein abundance, based on summed protein levels per class. For the analysis in Figure 3B and C protein abundance values were log2-transformed. Median expression and robust standard deviation (rSD) are used/indicated in Figure 3A, B and D. Abbrev.: OB: osteoblast, OC: osteoclast, LF: lung fibroblast, MEF: mouse embryonic fibroblast, MP: macrophages.

### Cell type specific dynamics of functionally connected lysosomal protein groups

While the previous analyses enable the identification of protein class-specific differences, they fail to reveal a possible heterogeneity within a respective group of proteins. We, therefore, further investigated the relative compositions of individual protein groups (Fig. 5). With respect to structural membrane proteins, we observed a similar behavior across cell lines for most proteins, with the exception of Lamp1/Lamp2 ratios. While MEFs expressed both proteins at similar ratios, in primary cells copy numbers of Lamp1 exceeded those of Lamp2 by more than 2-fold (Fig. 5A, Fig. S7D). As we identified the localization of Lamp1 to lysosomes to be well below 100% in MEFs (Fig 3D), and we are so-far not aware of non-vesicular localization of Lamp1 which was further reported to localize to other punctate structures than hydrolytic Ctsd-positive lysosomes [106], this finding is in line with either Lamp1 only positive lysosomes, or an entirely different compartment.

**Figure 5:**
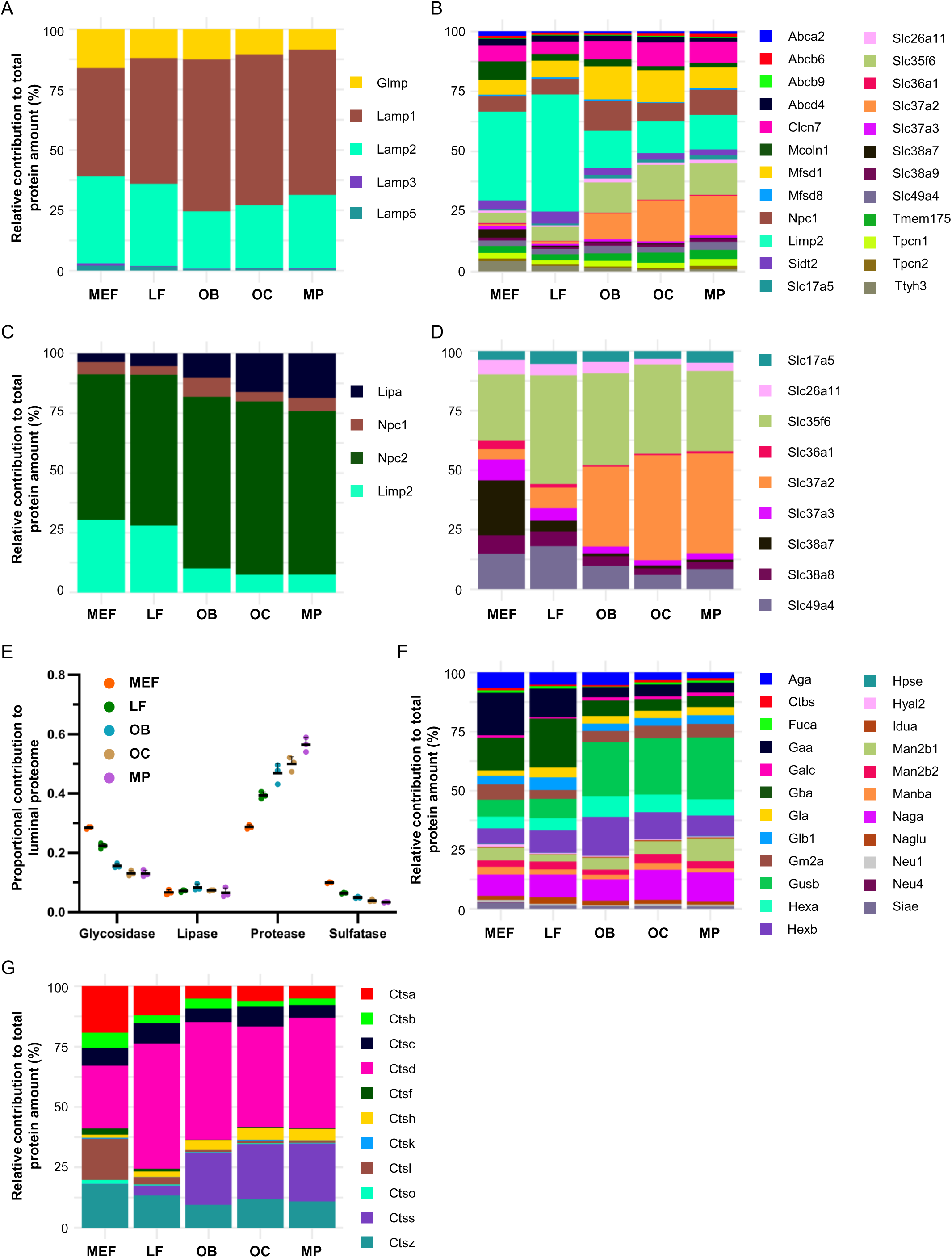
Quantitative composition of classes/categories in individual cell types. Shown is the relative contribution (%) as part of the total protein amount per class/category for A) structural membrane proteins, B) transporters, channels and exchangers, C) proteins involved in cholesterol homeostasis, D) solute carrier proteins, F) glycosidases, and G) cathepsins. E) shows the contribution of four hydrolase subtypes as a fraction of the total luminal proteome. Reported is the average across biological replicates (n=3); error bars indicate the standard deviation (SD). Abbrev.: OB: osteoblast, OC: osteoclast, LF: lung fibroblast, MEF: mouse embryonic fibroblast, MP: macrophages.

For the other major group of membrane proteins, which is transporters, channels, and exchangers, especially the cholesterol channel-containing protein Limp2 [129], which constitutes almost 50% of transporters in LFs, and the phosphate to glucose 6 phosphate antiporter Slc37a2, showed strong discrepancies between MEFs/LFs and OBs/OCs/MPs (Fig. 5B). Interestingly, when taking into account also the other proteins known to be involved in cholesterol homeostasis in the lysosome, the only one whose difference in expression (inversely) correlated with Limp2 was Lipa, which catalyzes the hydrolysis of cholesterol esters originating from low density lipoproteins (LDLs [129]) drawing a picture with either high Lipa or Limp2 levels (Fig. 5C). This could be related to similar levels of lysosomal cholesterol in all these cell lines, but different origins with a higher contribution of extracellular LDL derived cholesterol in OBs/OCs/MPs, and higher shuttling of cholesterol from the ER to the lysosome in MEFs and LFs. The most abundant protein with regard to cholesterol metabolism was in all cell lines Npc2 (Fig. 5C), which solubilizes free cholesterol in the lysosomal lumen, and ranged also with respect to total lysosomal protein abundance among the highest proteins (Table S3). With regard to the total abundance of SLC transporters, Slc37a2, Slc35f6, and Slc38a7 dominated this group in different ratios for the individual cell lines (Fig. 5D). In how far the phosphate to glucose 6 phosphate antiport by Slc37a2 is a major feature of lysosomes in these cells remains questionable, as this protein was the least lysosomal-localized SLC in MEFs (Fig. 3F) and was shown previously to be localized at the ER [130]. On the other hand, the nucleotide-sugar transporter Slc35f6 [131] was almost exclusively localized at lysosomes in MEFs, and could be involved in the starvation-induced salvage and cellular reutilization of ribosome-derived nucleosides [132]. Slc38a7 was only highly expressed in immortalized MEFs, which could be related to the fact that these cells are grown in culture for a long time, as this protein was identified as the major exporter of glutamine in cancer cells [111], and MEFs are typically cultured in medium which is supplemented with high levels of glutamine. For ABC transporters, we also observed a certain heterogeneity in protein expression levels, but by far not as pronounced as for SLC transporters (Fig S7E).

Finally, we investigated the dynamics of lysosomal hydrolases between the individual cell types, categorizing them based on the type of substrate group. With respect to the relative summed abundance of individual classes, the MEFs/LFs luminal proteome contained higher levels of glycosidases and sulfatases, and OBs, OCs, and MPs markedly higher amounts of peptidases, while lipases were present at similar levels in all cell types (Fig. 5E). However, within these individual classes, the relative expression patterns of individual proteins were rather similar, and only individual proteins represented outliers with respect to the class-specific patterns. With respect to glycosidases, for example, Gaa and Gba, which are involved in glycogen degradation [133] and removal of a glucose moiety from the membrane lipid glucocerebroside [134], respectively, were markedly higher abundant in MEFs and LFs (Fig. 5F). In OBs, OCs, and MPs, on the other hand, Gusb, which catalyzes the removal of the sugar glucuronic acid, a part of large polysaccharides called glycosaminoglycans (GAGs), was very prominent. For peptidases, Ctsl showed almost exclusive expression in MEFs, while Ctss was virtually absent in these cells but expressed in all primary cell lines, with especially high amounts in OBs, OCs, and MPs (Fig. 5G). Ctss has been associated mainly with ECM remodeling and bone/cartilage-related processes, with strong affection of these tissues in case of Ctss deficiency, which is in line with the observed increased expression in the bone-related cell types in our analyses [135].

### Transcriptional and post-transcriptional regulation of lysosomal luminal proteins

The observed heterogeneity at the protein level could be due to varying transcription rates in the individual cell lines or post-transcriptional processes. It has been shown previously, for example, that cathepsin L, K and B expression levels are transcriptionally regulated by various transcription factors, e.g. NF-Y, Sp1, Sp3, erythroblast transformation-specific family factors [136, 137] as well as Mitf and TFE3 [138]. To discriminate between these two major possibilities, we performed individual qPCR analyses for 51 lysosomal luminal proteins for one fibroblast cell line (LF), one bone derived cell line (OBs) and macrophages (MPs) and correlated them with the respective protein levels (Fig. 6). Interestingly, we observed very similar trends of relative transcript and protein levels between the three cell lines, indicating that the observed differences in protein expression are predominantly regulated on a transcriptional level (Fig. S8). This was, for example, the case for Ctsd, for which high transcript levels also resulted in high levels of proteins. We did observe, however, also the case that relative protein levels exceed those of their respective mRNAs (e.g. Tpp1, Ppt1, or Arsb) or that high transcript levels resulted in virtually no protein output, which was especially the case for Ctsb, Psap A/B/C/D or Ctsk.

**Figure 6:**
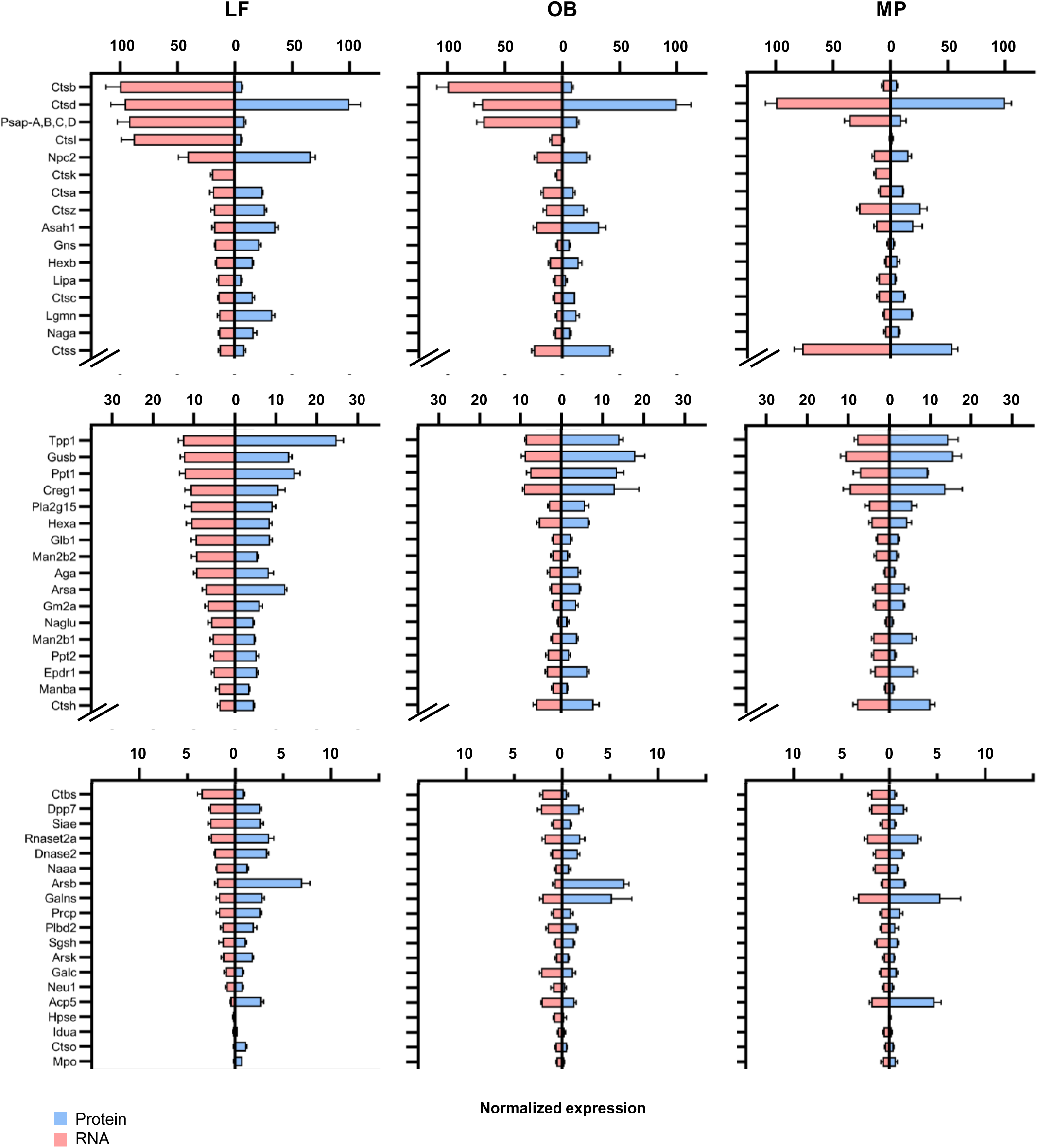
Comparison of RNA and protein expression profiles in primary cells. Mirror plots show normalized RNA and protein expression levels scaled to the dataset’s maximum expression. Error bars indicate the scaled standard deviation (SD). Abbrev.: LF: lung fibroblast, OB: osteoblast, MP: macrophages.

This could either be related to posttranscriptional regulation mechanisms, pronounced exocytosis of these proteins, or a high proteolytic turnover in the lysosome. The latter could be especially the case for Psap, as we also detected Psap B/D as the most abundant proteins in MEF-derived LEFs (Fig. S5).

## CONCLUSION

Our targeted MS platform combines MRM-MS with QconCAT-derived, absolutely quantified stable isotope-labeled standard peptides, enabling the absolute quantification of lysosomal proteins from whole-cell or tissue lysates. This robust, universally applicable strategy eliminates the need for lysosome enrichment and was used in this study for the absolute quantification of up to 143 lysosomal proteins across diverse cell types, including MEFs, LFs, OCs, OBs, and MPs. Based on the combination of MEF whole cell lysate and LEF data, we determined group-specific distribution patterns of lysosomal proteins, with a substantial proportion of membrane-bound proteins locating to subcellular compartments other than lysosomes. This suggests the existence of additional protein pools at other subcellular localizations, or the retention of lysosomal proteins outside this organelle. Furthermore, we identified non-lysosomal localization of luminal, hydrolytically active proteins, implicating the possibility of lysosomal hydrolase function and involvement in cellular processes beyond the lysosome itself. Across different cell types, we observed pronounced variations in the abundance of distinct lysosomal protein class levels and lysosomal proteome composition. This highlights the functional specialization of individual cell types, and hence their lysosomes. In general, primary cells exhibit higher abundances of hydrolases/luminal proteins, while the relative quantitative composition of the lysosomal membrane with respect to vATPase complex members and other lysosomal structural membrane proteins suggests to be preserved across cell lines. Of note, individual cell lines displayed unique expression dynamics for lysosomal hydrolases when distinguished by their catalyzed enzymatic reactions, thereby further revealing a lysosomal proteome tailored to their function. Within the protein groups, however, the individual composition was quite homogeneous, with only few selected proteins deviating strongly from the overall quantitative composition pattern. Based on the qPCR analysis of 51 lysosomal proteins, this cell type-specific protein expression appears to be, with few exceptions, tightly and directly regulated on the transcriptional level.

## Acknowledgement

Sandra Pohl received funding from the German Research Foundation (Deutsche Forschungsgemeinschaft, DFG) - grant PO 1539/1. Dominic Winter received funding from the DFG as part of the FOR2625. Sofía Fajardo-Callejon received funding from the German Academic Exchange Service (Deutscher Akademischer Austauschdienst, DAAD).

## Author contributions

Peter Robert Mosen: Data curation, Formal analysis, Software, Visualization, Writing – review & editing, Investigation, Validation, Methodology, Project administration, Writing – original draft; Biswajit Moharana: Data curation, Investigation; Sofía Fajardo-Callejon: Data curation, Formal analysis, Visualization, Investigation; Validation; Edgar Kade: Investigation; Norbert Rösel: Investigation; Maryam Omidi: Data curation, Investigation; Roman Sakson: Data curation, Formal analysis, Investigation; Thomas Ruppert: Resource; Sandra Pohl: Formal analysis, Resource; Volkmar Gieselmann: Funding; Dominic Winter: Methodology, Writing – review & editing, Project administration, Conceptualization, Resource; Supervision, Validation, Funding, Writing – original draft.

---

**Figure S1:**
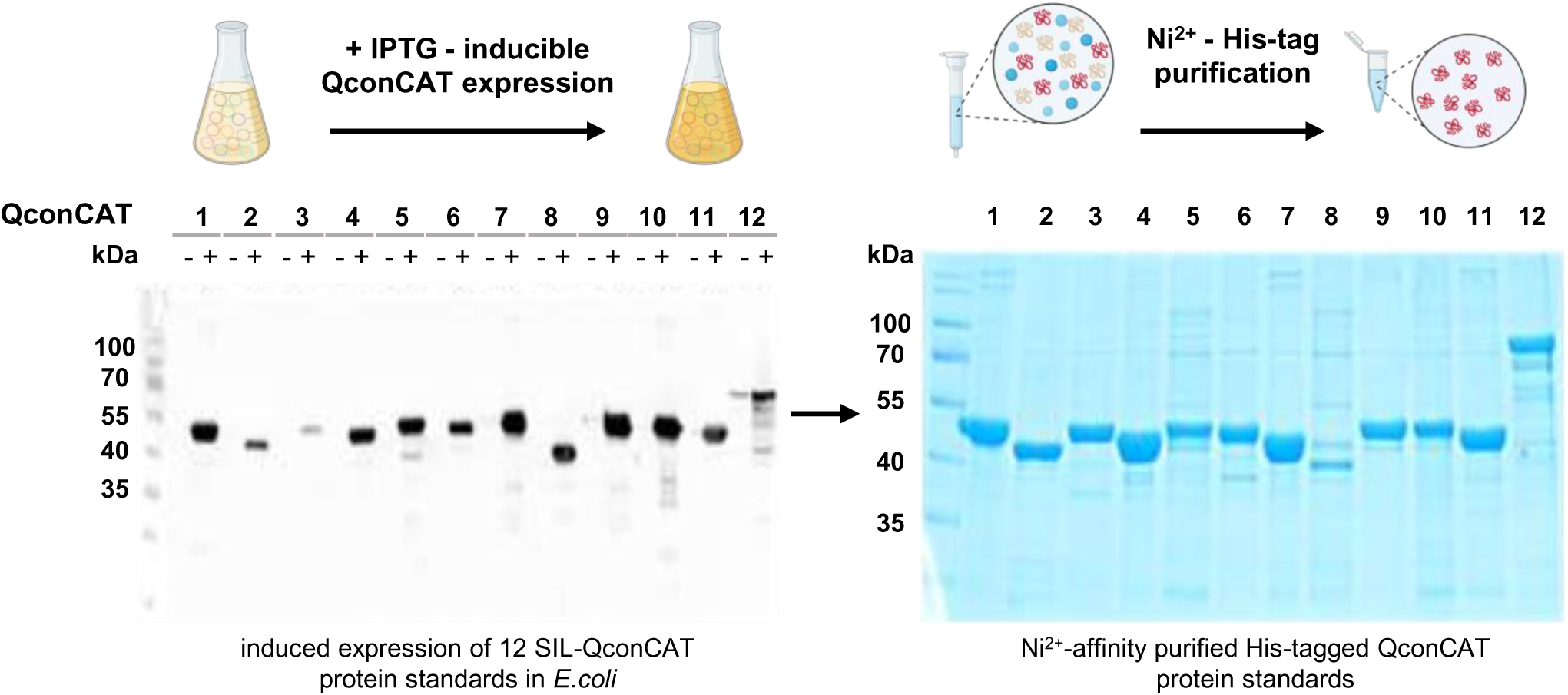
Expression and purification of lysosomal QconCAT protein constructs. **Expression of** 12 different QconCATs upon IPTG induction (+) was confirmed by Western Blot using a His-tag antibody. (-) shows non-induced controls. Heavy, stable isotope-labeled QconCATs were purified by Ni_2+-_affinity purification. Abbrev.: SIL: stable isotope labeled. Figure S1 was generated using BioRender.

**Figure S2:**
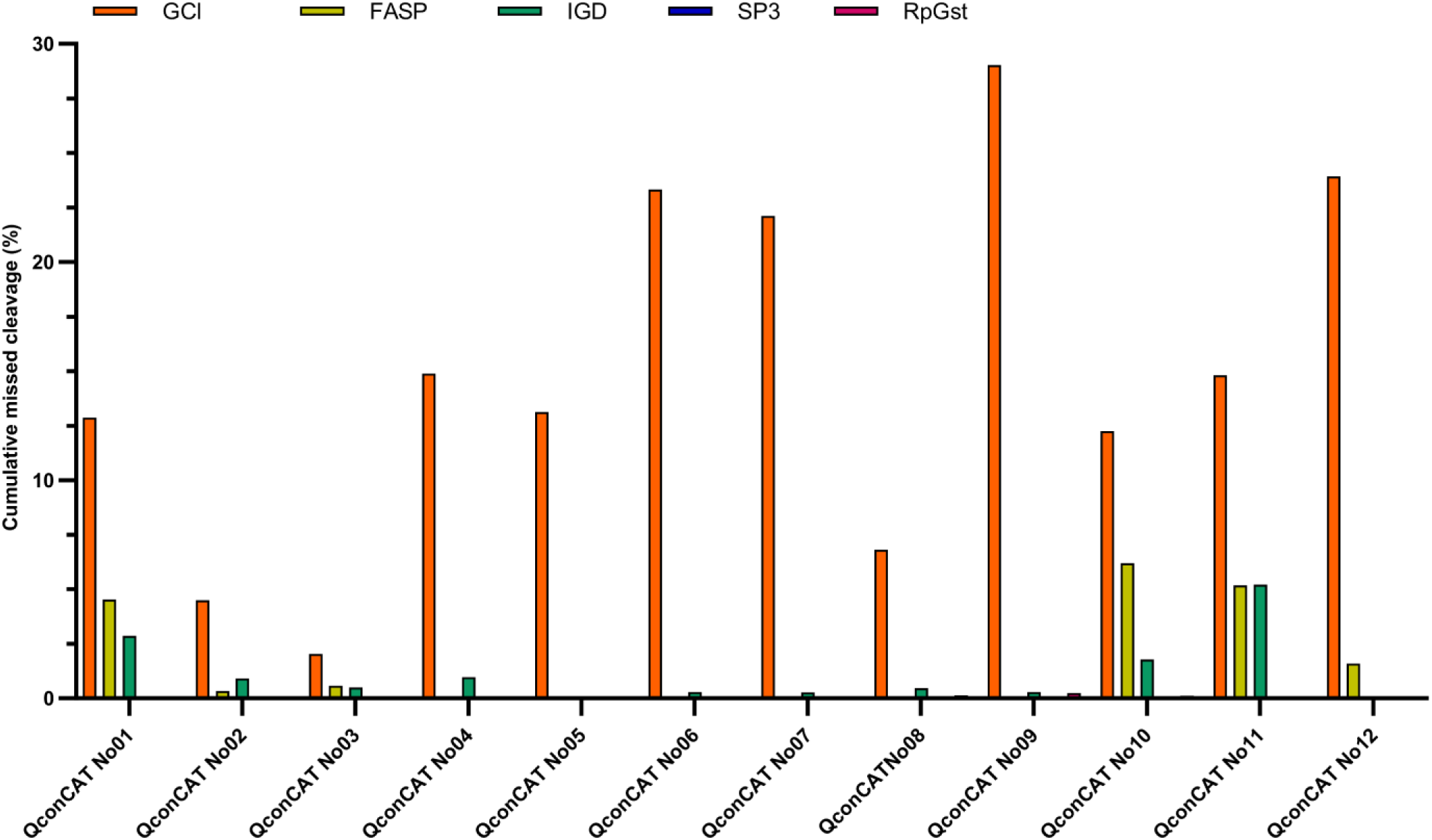
Evaluation and comparison of missed cleavage rates for individual protein standards. Expressed and purified QconCATs were digested using different digestion strategies. Missed cleavage rates reported represent the protein construct’s cumulative missed cleavage rate. Abbrev.: IGD: in gel digestion, FASP: Filter-Aided Sample Preparation, GCl: guanidinium chloride, RpGst: RapiGest, SP3: solid-phase enhanced sample-preparation

**Figure S3:**
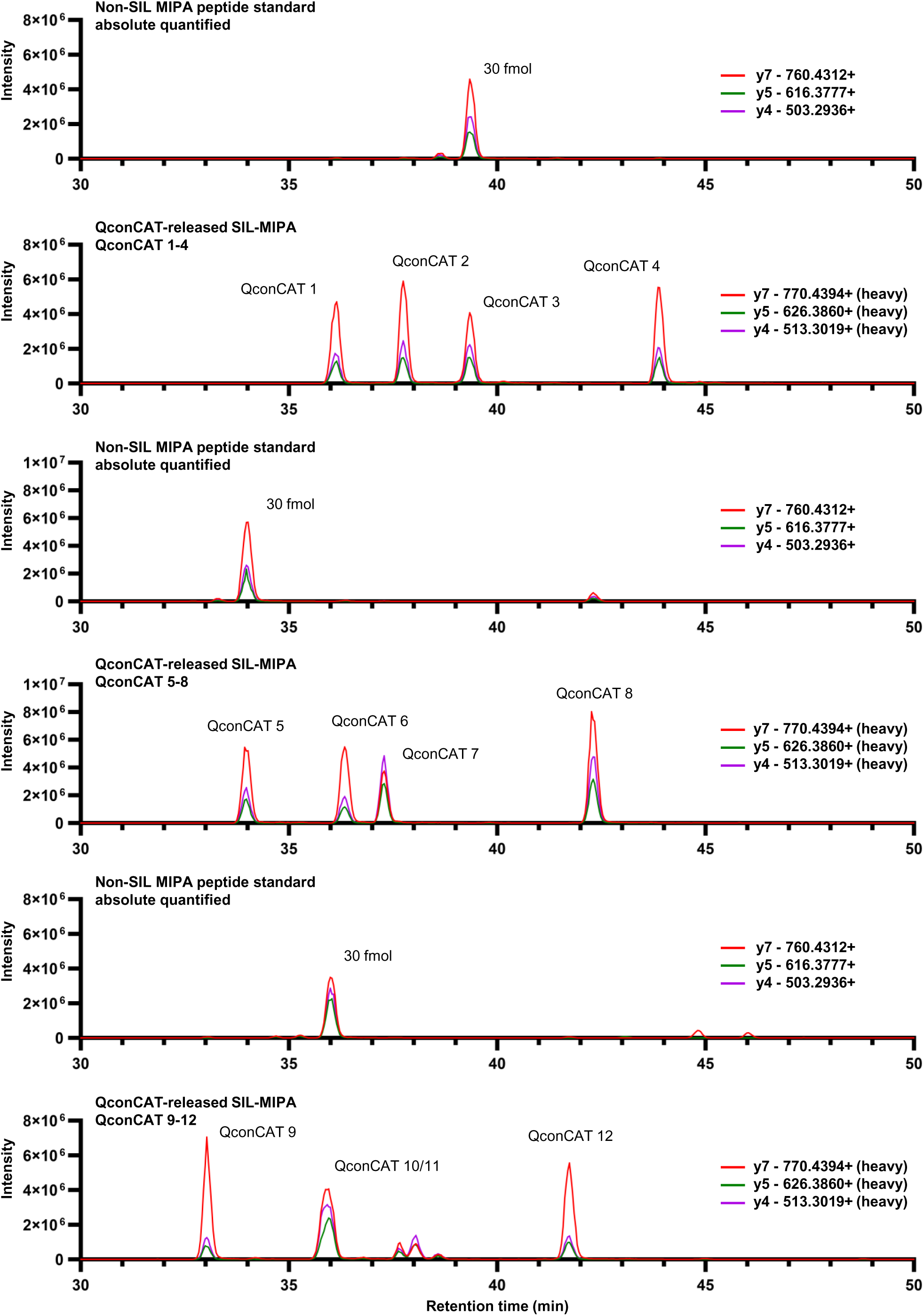
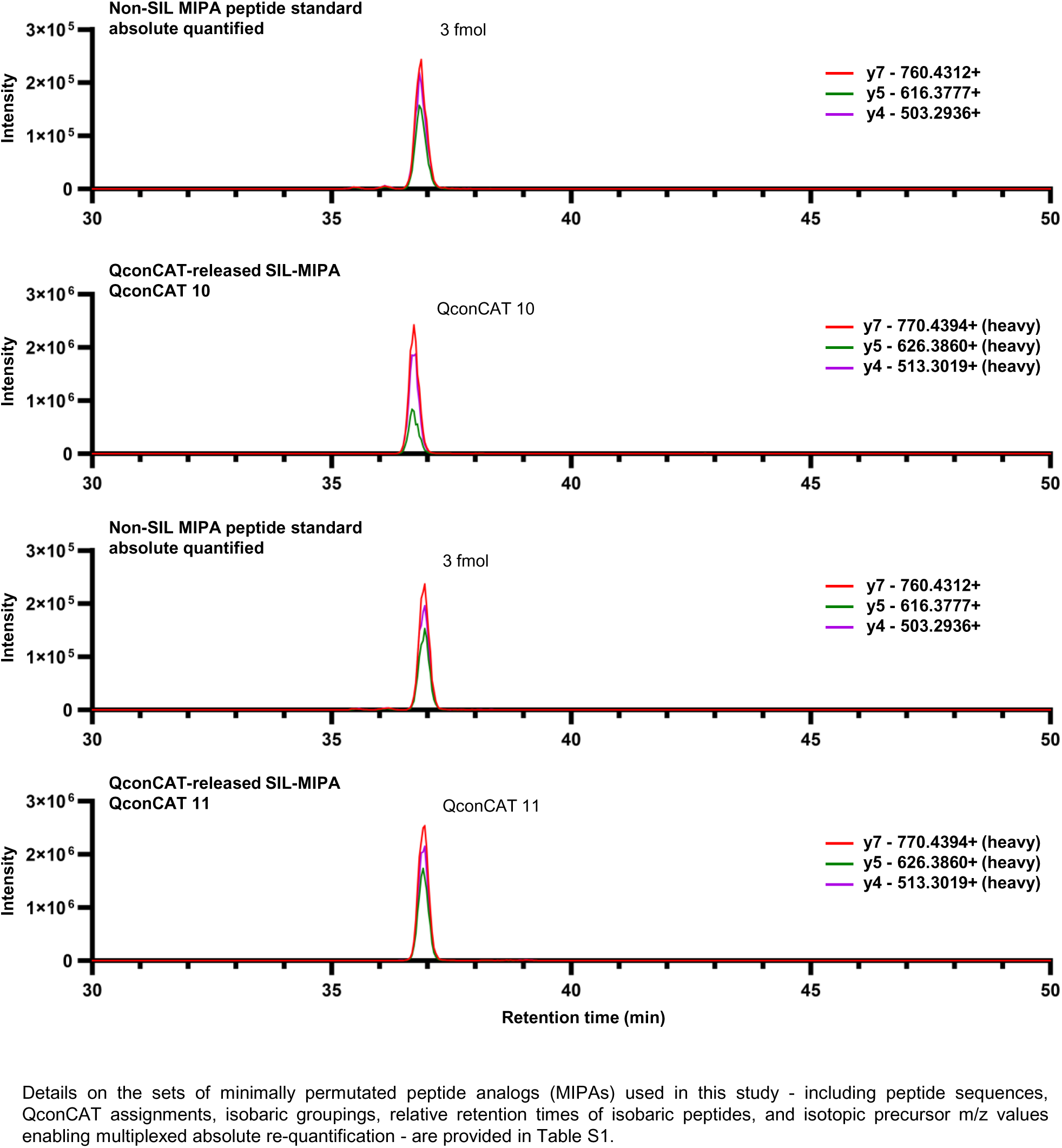
Re-quantification of 12 stable isotope-labeled QconCAT peptide standards. Sets of four SIL protein/peptide standards (heavy) were re-quantified against a known amount of non-SIL peptide standard (light). MIPA peptides representative of QconCAT 10 and 11 co-elute, and had to be re-quantified separately (Continuation Figure S3). Abbrev.: SIL: stable isotope labeled, MIPA: minimally permutated peptide analog.

**Figure S4:**
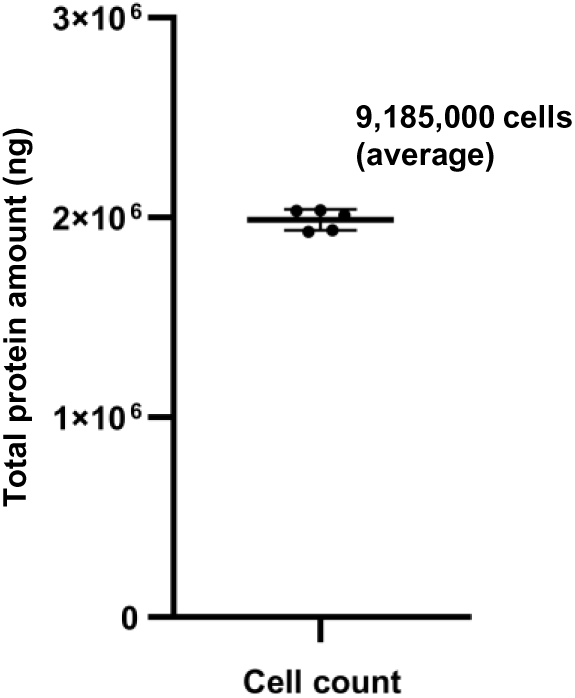
Quantification of protein content in mouse embryonic fibroblast cells. For defined cell numbers, the protein content was determined after lysis. Shown is the average (n=5), error bars indicate the standard deviation.

**Figure S5:**
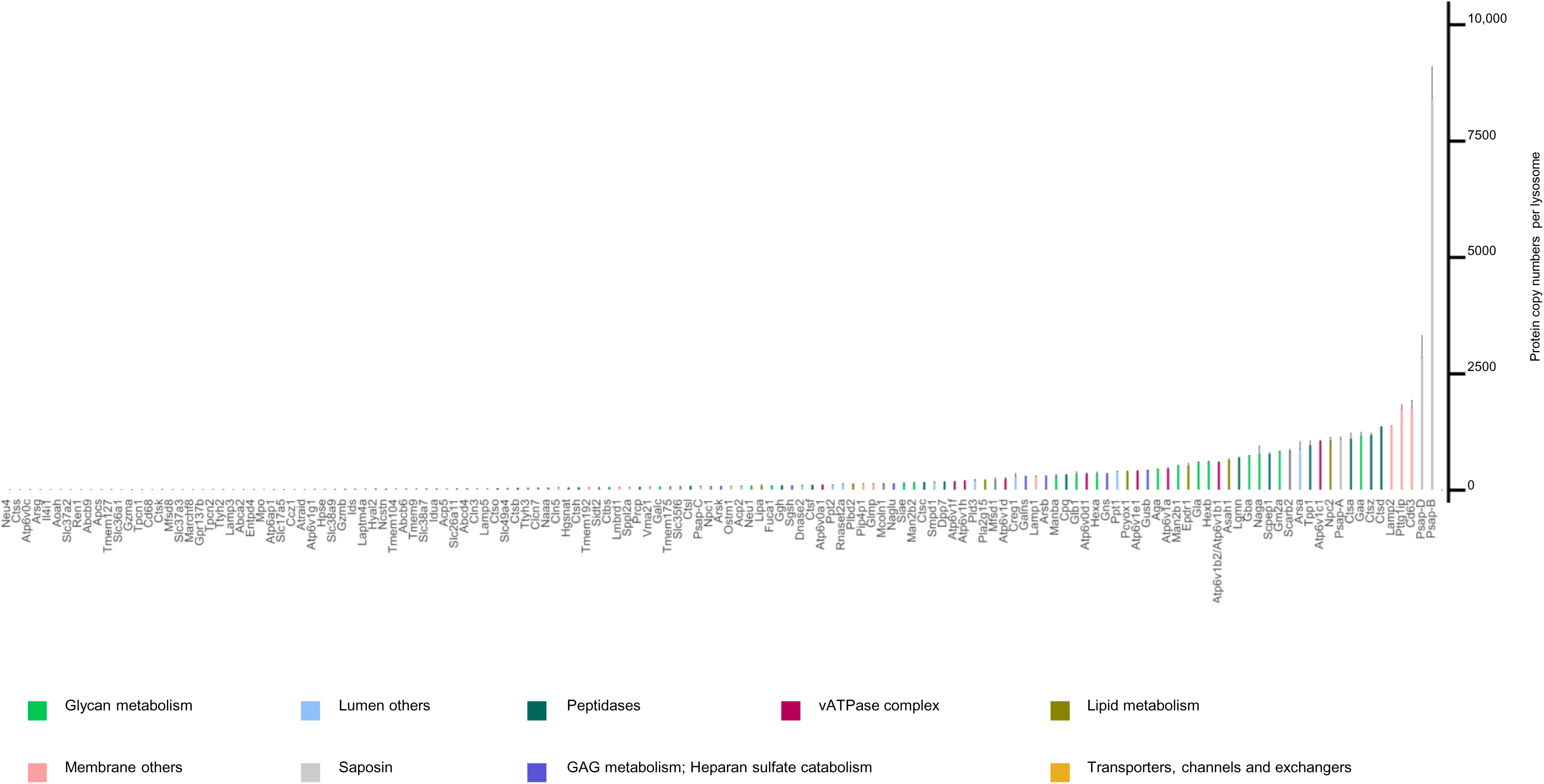
Lysosome-specific protein copy number estimation. Protein copy numbers for 141 proteins determined from LEFs are shown. Proteins are associated with nine different lysosomal classes/categories. Reported are median values across biological replicates (n=3); error bars indicate the robust standard deviation (rSD). Abbrev.: LEF: lysosome-enriched fraction.

**Figure S6:**
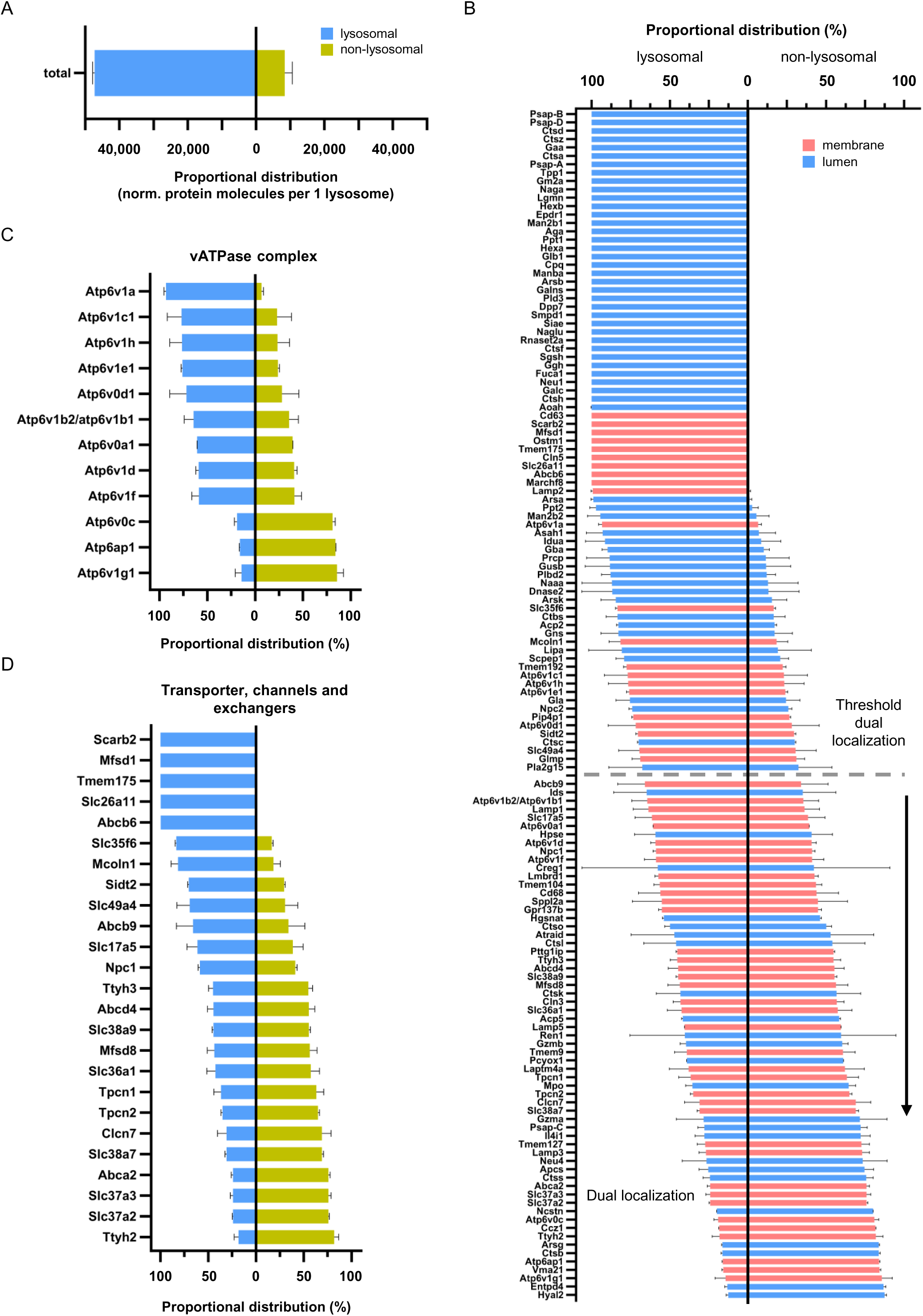
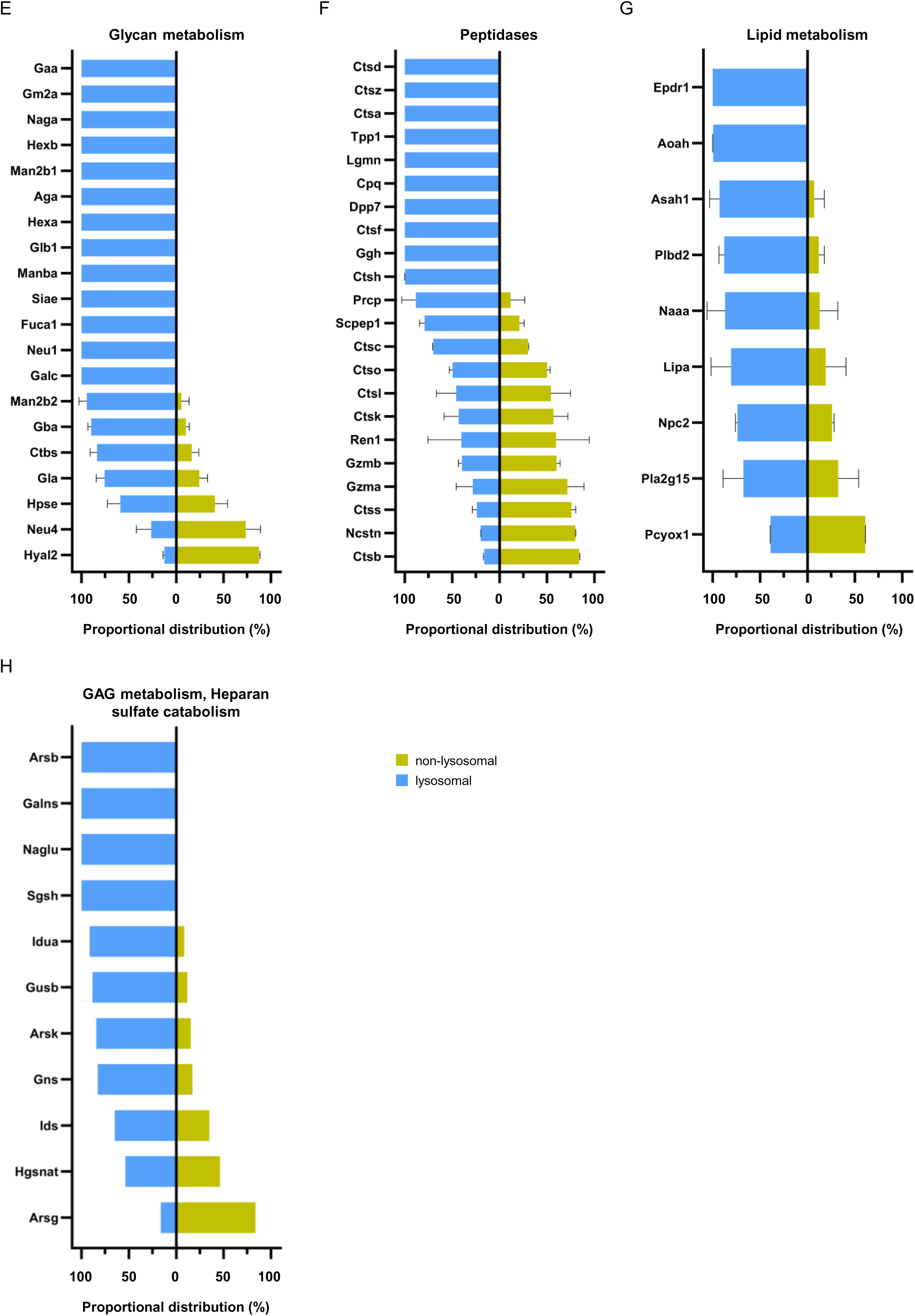
**Localization study for lysosomal proteins**. A-H) Proportional distribution of lysosomal protein molecules located in/at the lysosome (lysosomal, left) and outside the lysosome (non-lysosomal, right). B) Shown is the proportional distribution for the sum of protein molecules. C) Each horizontal bar represents a protein. Colors indicate the topology. The grey horizontal line indicates the threshold cut-off for dual localization. In this analysis, we observed for several proteins values exceeding 100%. As it is not possible, that more of a certain protein is present at lysosomes than in the whole cell, we argued that this finding is probably due to a stronger suppression of individual peptide signals in whole cell lysates, and hence a slight underestimation of their abundance. We, therefore, defined these proteins as fully lysosome-localized (see method section). Median values across biological replicates (n=3) are reported; error bars indicate the robust standard deviation (rSD).

**Figure S7:**
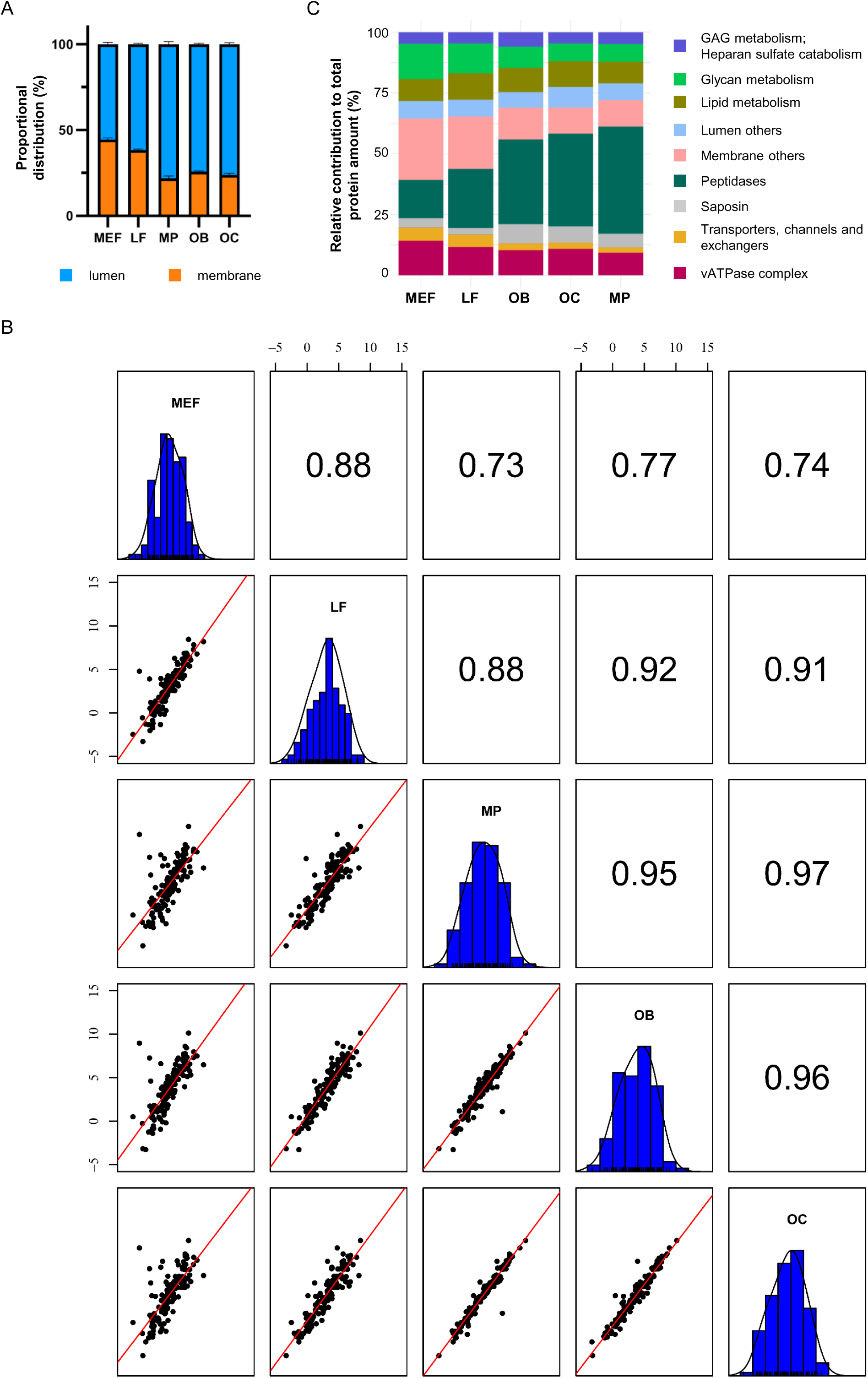
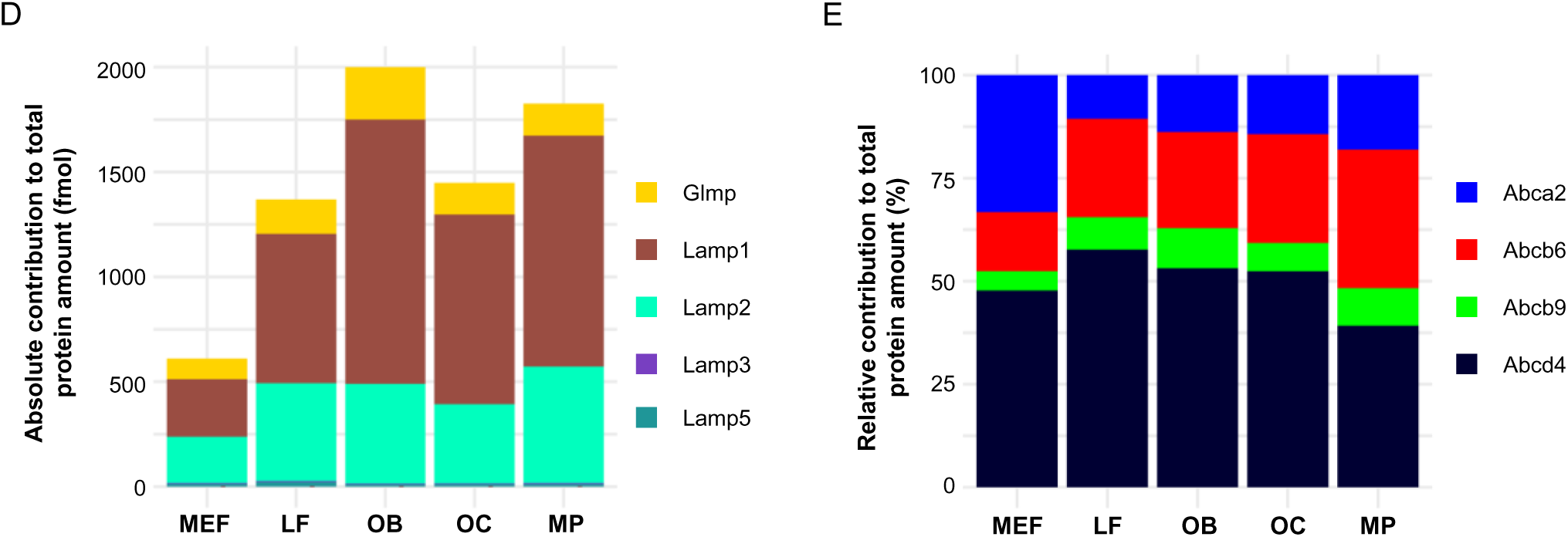
Lysosomal protein expression analysis for individual cell types. A, C, D and E) Shown is the relative (%) and absolute (fmol) contribution of individual protein amounts/classes to the total protein amount for A) luminal and membrane protein localization, C) nine lysosomal functional and structural classes, D) lysosome-associated membrane proteins, and E) ATP-binding cassette transporters. B) Correlation of lysosomal protein expression profiles. The Pearson correlation of log2- transformed absolute protein amounts was calculated. Abbrev.: OB: osteoblast, OC: osteoclast, LF: lung fibroblast, MP: macrophages.

**Figure S8:**
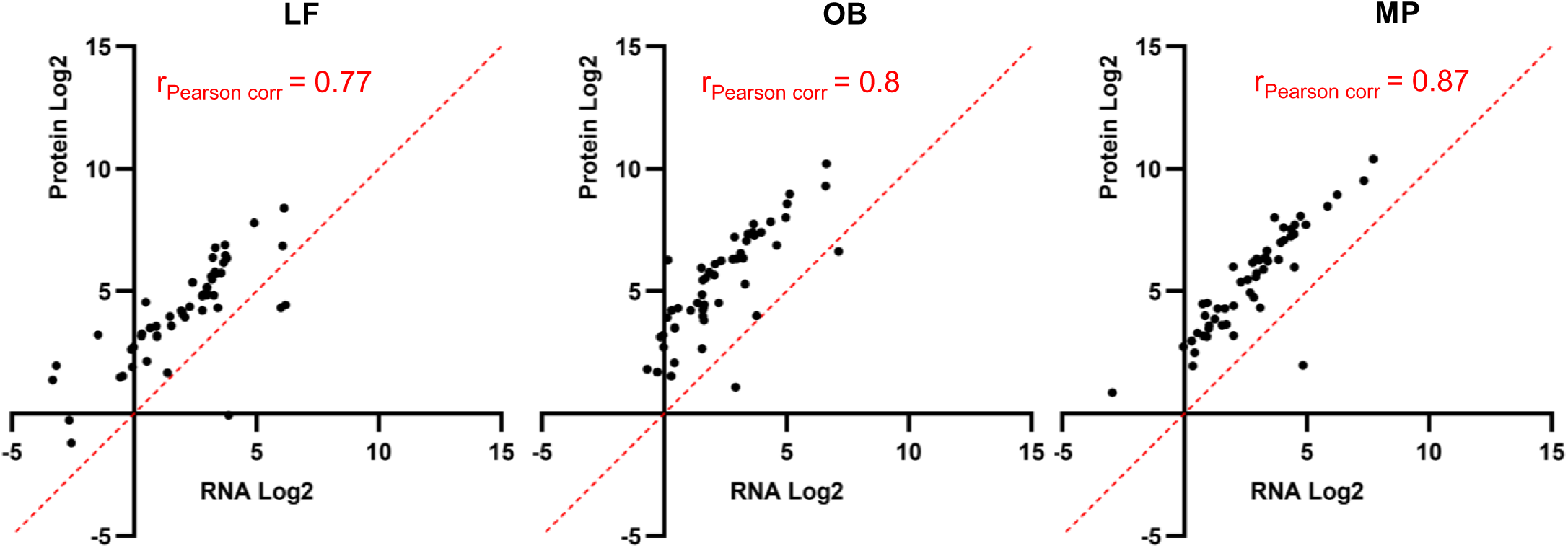
Correlation of RNA and protein expression levels between primary cell types. Expression values are Log2-transformed, the Pearson correlation is reported. Abbrev.: LF: lung fibroblast, OB: osteoblast, MP: macrophages.

